# Adaptive introgression between two *Drosophila* species enhances heat tolerance despite barriers to gene flow

**DOI:** 10.64898/2026.01.13.699320

**Authors:** Noora Poikela, Rhonda R. Snook, Jonna Kulmuni, Michael G. Ritchie

## Abstract

As global temperatures rise and heat waves become more frequent, populations must adapt rapidly to avoid local extinctions. Adaptive introgression allows species to quickly acquire adaptive genetic material from close relatives. However, certain genomic regions, containing barrier loci, may resist introgression. We experimentally tested whether strong heat stress selection can overcome such barriers and facilitate the transfer of adaptive alleles to improve heat tolerance. We also examined whether chromosomal inversions, which can tightly link adaptive and barrier loci through reduced recombination, promote or prevent introgression. We conducted a hybridize, evolve, and re-sequence experiment by crossing the heat-sensitive *D. flavomontana* with the more heat-tolerant *D. montana*, exposing heat-selection lines to transient heat for three generations while maintaining control lines. We also mapped long-term barriers to gene flow between natural populations using a demographically explicit genome scan. The heat-selection lines exhibited approximately twice the introgression and higher male fertility under heat stress compared to control lines and the heat-sensitive parental *D. flavomontana*. This introgression was primarily located in colinear autosomal regions and correlated with improved heat tolerance. Some introgression occurred also in inverted autosomal regions in the heat-selection lines but not in the control lines. The X chromosome, with three overlapping inversions, resisted introgression in both heat-selection and control lines, thereby maintaining a strong species barrier. Finally, genetic barriers in the control lines significantly overlapped with long-term barriers, but this overlap was less pronounced in the heat-selection lines. Our study demonstrates that strong selection can lead to adaptive introgression despite long-standing genetic barriers. These findings highlight that introgression can serve as a rapid source of adaptive genetic material under stressful thermal conditions, potentially mitigating climate-induced population extinctions.

## Introduction

As global temperatures rise and heat waves become more frequent [1], populations must adapt rapidly to these changes to avoid local extinctions. Although new mutations and existing genetic variation provide essential material for natural selection, both sources have limitations [2]. Populations may acquire new, adaptive genetic material from closely related species through hybridization and interspecific gene flow, a process known as “adaptive introgression” [2–7]. This process is particularly relevant as climate change induces species redistribution, bringing previously isolated related species into secondary contact [8]. However, selection may act against introgression at specific genomic regions, known as barrier loci, to mitigate any negative effects of hybridization and resulting gene flow [9–11]. The extent to which adaptive introgression between species can overcome strong barriers in empirical systems remains understudied (but see [12–14]). Here, we experimentally test if strong heat stress selection can overcome interspecific barriers and improve heat tolerance during the first few hybrid generations in species despite strong reproductive isolation.

Recent studies emphasize that species distributions can be shaped by thermal limits to reproduction rather than to viability [15,16]. High temperatures are particularly harmful to male fertility, reducing sperm production, viability, and insemination success, and compromise the reproductive potential and lifespan of offspring sired by males exposed to heat [17]. This fertility reduction could limit where species can live, making it a significant driver of biodiversity declines due to global warming. In *Drosophila*, both brief heat shock and experimental evolution under increasing temperatures found that male heat-induced sterility temperatures were better predictors of species distributions than survival limits [15,18]. Moreover, extinction risk was predicted by male sterility temperature [18]. While populations can exhibit variation in resilience to heat-induced male sterility [19], genetic variation is limited [20,21] and, under thermal experimental evolution, species did not shift the temperature that induced male sterility [18]. Thus, standing genetic variation may be insufficient for rapidly evolving resilience to male heat-induced sterility during strong thermal selection. Adaptive introgression could be one mechanism that provides a way out of this problem [7].

However, predicting the outcomes of introgression is difficult. Although it can be a crucial mechanism for evolutionary rescue, genetic barriers may counteract introgression [9–11]. Barrier loci are those in which gene exchange between lineages is selected against. They may contribute to local adaptation, assortative mating, differences in the number of deleterious variants carried by the hybridizing species (i.e. hybridization load), or reduced hybrid fertility or viability due to genetic incompatibilities arising from negative epistatic interactions (Bateson-Dobzhansky-Muller incompatibilities, BDMIs [22,23]) [24–26]. Barriers to gene flow can be detected by studying gene exchange patterns in hybrids within natural hybrid zones or hybrid populations [27–30] and through laboratory crosses [31–34]. Furthermore, recent advances in demographic modeling have enabled the detection of genomic barriers to gene flow throughout species’ evolutionary history, indicated by genomic regions with reduced effective inferred migration rates (*m_e_*) [35,36]. Here, we define barriers observed in laboratory crosses as “contemporary barriers”, while those identified through demographic modeling are referred to as “long-term barriers”. A few studies have examined the overlap of barriers detected using different approaches, allowing fo r reliable barrier mapping across different time scales [29,30,37]. However, the extent to which strong positive selection can overcome the costs associated with barriers remain poorly understood.

The effect of barrier loci can extend to linked neutral and adaptive genomic regions, creating areas of reduced introgression, particularly in regions of low recombination [11]. For example, chromosomal inversions, regions of the genome where the gene order is reversed relative to the standard form, suppress recombination across the inverted sequence due to disrupted alignment of homologous chromosomes [38,39]. This suppression facilitates the accumulation and preservation of alleles associated with local adaptation and the build-up of barriers to gene flow, keeping them tightly linked [40–45]. Consequently, inversions can either promote or hinder adaptive introgression, depending on the extent to which they contain adaptive or barrier loci, and the relative strength of selection. If inversions harbor strong barriers to gene flow, the entire inversion may resist introgression since recombination cannot separate adaptive and neutral alleles from linked maladaptive alleles. For example, introgression was less likely to persist in regions of low recombination in natural hybrid populations of two swordtail fish species [27]. Conversely, if strong positive selection outweighs the costs of maladaptive loci, the entire inverted region may be transferred as “adaptive cassette” from one species to another through introgression [46–49].

Two species of the *Drosophila virilis* group, *D. flavomontana* and *D. montana*, offer a valuable system to study the potential of introgression in adaptation to warming temperatures and the interplay between adaptive introgression, barrier loci, and recombination-suppressing inversions. *D. flavomontana* shows one of the largest temperature differences between thermal fertility limits and survival limits among *Drosophila* species [15]. In other words, males become sterile at much lower temperatures than those that are lethal. Consequently, the distribution of *D. flavomontana* is predicted to reduce 42-63% by 2080 due to increasing temperatures and the sterilizing effect of high temperatures on males [15]. In contrast, *D. montana* shows a much smaller difference between temperatures that sterilize males and those that kill them, and it maintains fertility at higher temperatures than *D. flavomontana* [15]. Therefore, introgression from *D. montana* could aid *D. flavomontana* in adapting to heat stress. Despite diverging approximately 2.5 million years ago, possessing alternatively fixed inversions that resist gene flow [34,50], and exhibiting strong reproductive isolation [51], these species can still interbreed in the laboratory [34]. Specifically, *D. flavomontana* females can be crossed with *D. montana* males, and the resulting F_1_ females can be backcrossed with either parental species, though F_1_ males are sterile [34,51,52]. Furthermore, interspecific hybrids have reportedly been found in nature [52,53].

In this study, we aim to assess the extent and pattern of introgression during the early generations of hybridization between species with strong reproductive isolation, and ask the following questions: 1) Does strong heat selection facilitate the transfer of adaptive alleles from the heat-tolerant *D. montana* to the heat-sensitive *D. flavomontana* through introgression, thereby improving heat tolerance (measured as heat-induced sterility) despite barriers to gene flow? 2) Do alternatively fixed chromosomal inversions facilitate or hinder adaptive introgression? 3) Does heat stress selection overcome contemporary barriers, and do these barriers coincide with long-term barriers to gene flow? To address these questions, we conducted a hybridize, evolve, and re- sequence (HER) experiment by crossing *D. flavomontana* females with *D. montana* males and backcrossing the F_1_ females with *D. flavomontana* males (Fig. 1). The resulting BC_1_ hybrids (with 25% *D. montana* ancestry) were divided into two groups: one exposed to transient heat stress selection for three generations (heat-selection treatment) and the other kept as controls (control treatment). After the selection, both groups were phenotyped for heat-induced sterility and pool-sequenced to compute hybrid indices (HI; i.e. amount of introgression from *D. montana* into *D. flavomontana*). Additionally, we mapped long-term barriers between natural populations of these species using a demographically explicit genome scan (gIMble; [36]). We hypothesize that strong heat stress selection will counteract genetic barriers, allowing heat-adapted alleles from the heat- tolerant species to transfer to the heat-sensitive species and improve heat tolerance. We also expect alternatively fixed inversions between the species − one each on autosomes 4 and 5, and three overlapping inversions on the X chromosome (Table S1) [50] − to harbor adaptive alleles which could act as adaptive cassettes transferred between the species. However, if adaptive alleles are tightly linked to strong barrier loci due to reduced recombination within these inversions, they may resist introgression.

**Figure 1.**
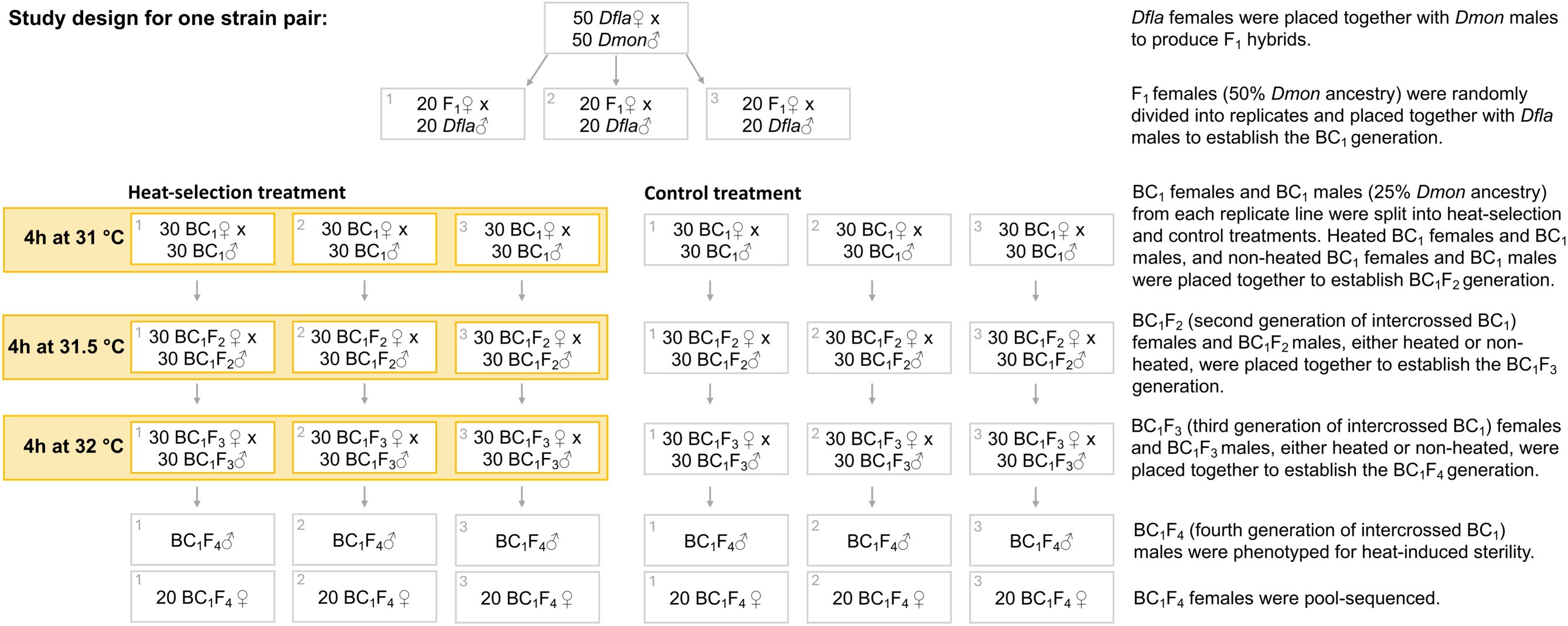
Outline of the hybridize, evolve, and re-sequence (HER) experiment illustrated for one strain pair (the experiment included two strain pairs, i.e. six heat-selection and six control lines altogether). The HER experiment was initiated by crossing *D. flavomontana* females with *D. montana* males. The resulting F_1_ females were divided into replicates and backcrossed with *D. flavomontana* males (replicate numbers shown in the upper left corner of the boxes). In the first backcross (BC_1_) generation, replicate lines were split into heat-selection and control treatments. Heat selection began with the BC_1_ generation, where flies were subjected to four hours at 31°C. Hybrids of the next two generations (BC_1_F_2_ and BC_1_F_3_) were exposed to 31.5 °C and 32 °C for four hours, respectively. The controls were not exposed to heat but were otherwise handled identically. In the BC_1_F_4_ generation, the heat-selection and control lines were sequenced and phenotyped. Further details in Materials and Methods.

## Results

### Heat stress induces greater sterility in D. flavomontana males compared to D. montana males

Before conducting the hybridize, evolve, and-re-sequence (HER) experiment, we tested what temperatures induce sterility in *D. flavomontana* males, and whether *D. montana* is more tolerant of the extreme temperatures. The fertility of *D. flavomontana* males decreased with increasing temperatures (Fig. 2A; GLM: z_27_ = -5.56, P < 0.0001), being as low as 27% at 32 °C. Although heat also induced sterility in *D. montana* males, they exhibited higher fertility compared to *D. flavomontana* males at both 31 °C and 32 °C (GLM: 31 °C: z_1,8_ = 5.50, P < 0.0001; 32 °C: z_1,7_ = 5.07, P < 0.0001; Fig. 2A). In contrast, when both females and males of each species were exposed to the control temperature of 20 °C, or when females of both species were exposed to temperatures between 28°C and 32 °C, their fertility remained around 90% on average (Fig. S1). In summary, temperatures of 31-32 °C cause 50-80% sterility in *D. flavomontana* males but have weaker effects on *D. montana* males. Therefore, these temperatures were used in the HER experiment (outlined in Fig. 1).

**Figure 2.**
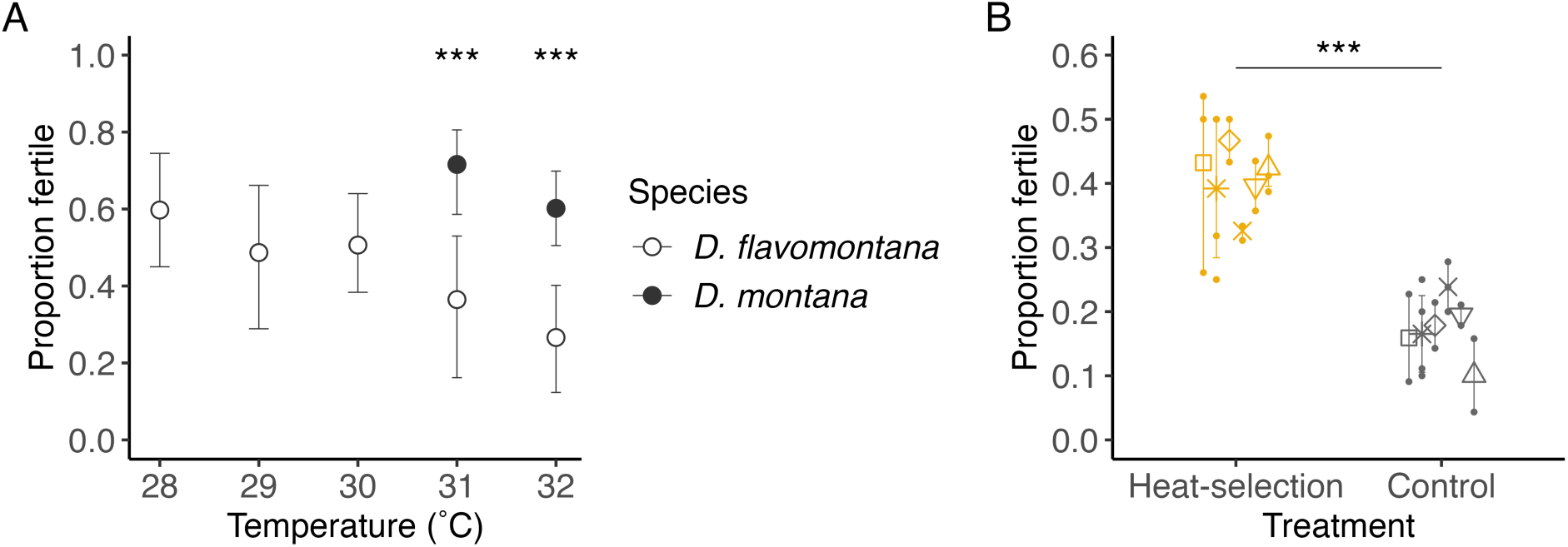
The effect of heat on male fertility between *D. flavomontana* and *D. montana*, and between the control and heat-selection treatments. (A) Proportion of fertile *D. flavomontana* and *D. montana* males after heat exposure. *D. flavomontana* males were exposed to five different temperatures, and *D. montana* males to two temperatures for four hours to estimate the effects of heat on male fertility in both species. Black stars indicate significant differences between the *D. flavomontana* and *D. montana* males at 31 °C and 32 °C (GLM: ***P<0.001). (B) Proportion of fertile males in the six control and heat-selection lines after exposing them to 32 °C for four hours. Black stars indicate significant differences between the control and heat-selection lines (GLMM: ***P<0.001). A higher proportion of fertile males indicates better tolerance to warm temperatures. Error bars represent 95% confidence intervals. The data underlying this figure can be found at doi.org/10.5281/zenodo.22277897.

### Heat stress selection increases male fertility thermal limits through introgression of heat-adapted alleles

After the HER experiment (Fig. 1), we assessed whether the heat-selection lines exhibited improved fertility compared to the control lines, and whether this improvement was accompanied by greater introgression from the more heat-tolerant *D. montana*. For this, we measured the male fertility of the control and heat-selection lines by exposing them to 32 °C for four hours. The heat- selection lines showed, on average, 2.2 times greater male fertility compared to the control lines (40% vs. 17%; GLMM: z_1,28_ = 6.37, P < 0.0001; Fig. 2B), indicating that the heat stress selection improved male ability to tolerate extreme temperatures. When compared to the parental species at 32 °C, the mean fertility of the heat-selection lines (40%) was higher than that of parental *D. flavomontana* (27%) but lower than that of parental *D. montana* (60%; Fig. 2). The mean fertility of the control lines was the lowest of all (17%).

A total of 700,751 diagnostic SNPs − SNPs that are differentially fixed between the species based on both pooled parental samples and samples sequenced from natural populations − were used to calculate hybrid indices (HIs) in the control and heat-selection lines (Table S2). The hybrid index indicates the amount of introgression from the heat-tolerant *D. montana* into the heat-sensitive *D. flavomontana*. Across all genomic regions, the mean HI of the heat-selection lines was nearly twice as high as that of the control lines (5.6% vs. 2.9%; Fig. 3, 4A, S2; Table S2). This indicates a higher overall introgression from the more heat-tolerant *D. montana* in the heat-selection compared to the control treatment.

**Figure 3.**
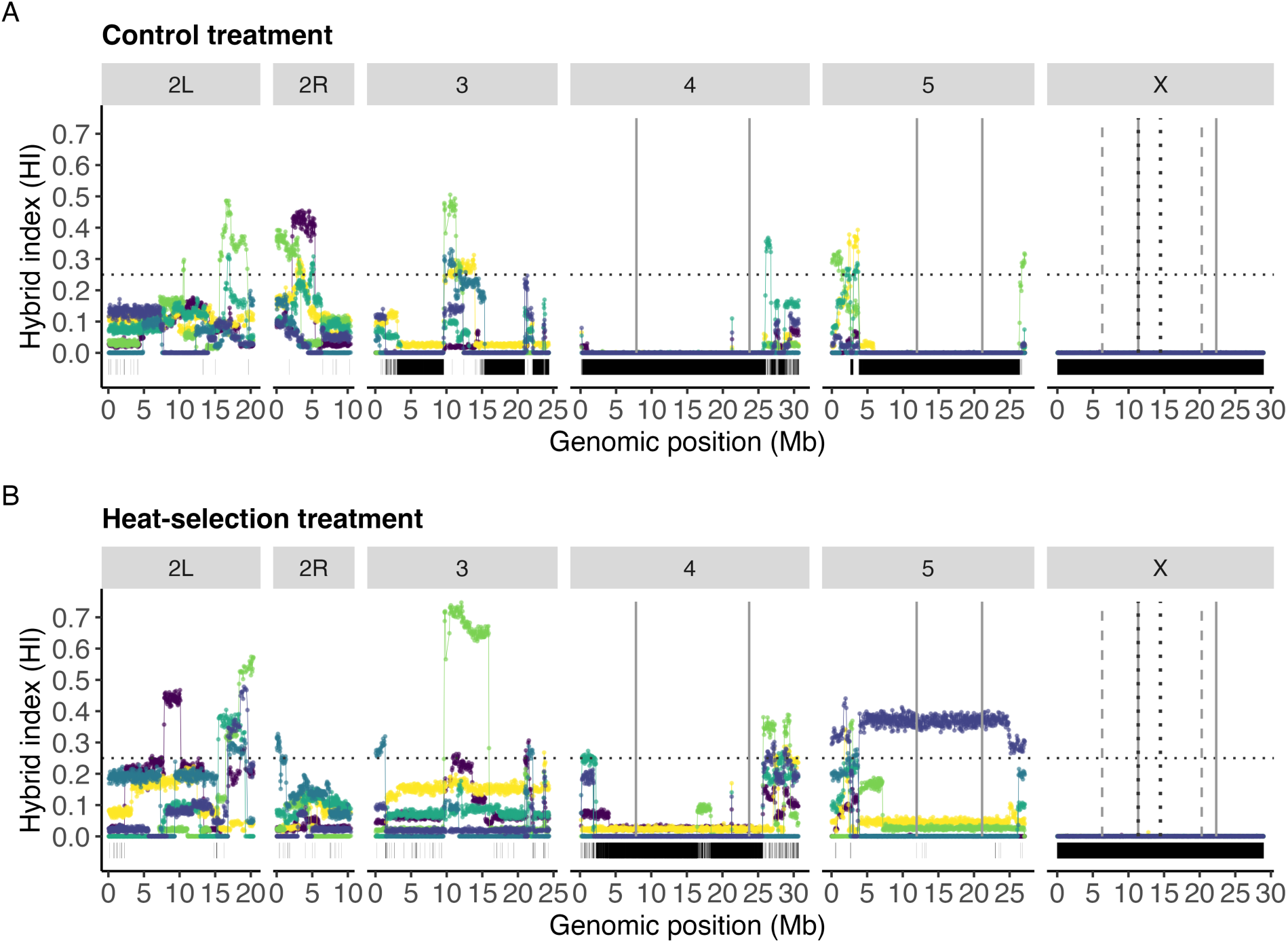
Genome-wide variation in hybrid index (HI) in the control and heat-selection treatments. HI was plotted in windows of 200 non-overlapping SNPs along the genome in (A) the control and (B) the heat selection lines (relative differences between the lines shown in Fig. S2A). Different colors indicate different replicate lines. The dashed horizontal line indicates the expected *D. montana* ancestry at the start of the hybridize, evolve, and re-sequence (HER) experiment. Vertical solid and dashed lines denote the breakpoints of the alternatively fixed chromosomal inversions between *D. flavomontana* and *D. montana*. Black segments at the bottom of the plots represent SNPs with an HI of 0, indicating no introgression from *D. montana* in any of the six control or six heat-selection lines (strong contemporary barriers). Note that the size of a segment is not proportional to a single SNP. These data are summarized in Fig. 4 and Table S2. The data underlying this figure can be found at doi.org/10.5281/zenodo.22277897.

**Figure 4.**
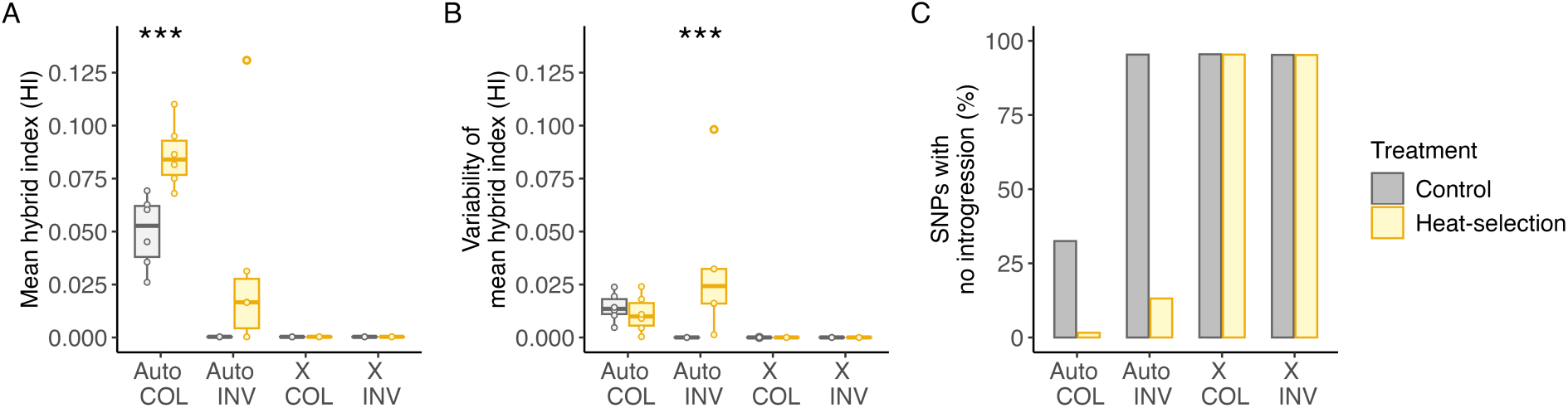
Comparison of introgression and strong contemporary barriers between the control and heat- selection treatments. (A) Mean hybrid index (HI), representing the amount of introgression from the more heat-tolerant *D. montana*, between the control and heat-selection treatments. Differences between the respective heat-selection and control replicate lines are shown in Fig. S2B. (B) Variability in mean HI, calculated as the absolute deviation from the mean, between the control and heat-selection treatment. (C) Proportion of SNPs with no introgression from *D. montana* (HI = 0) in any of the six control or six heat- selection lines, indicating strong contemporary barriers. Each metric was calculated separately for colinear (COL) and inverted (INV) autosomal (Auto) and X chromosomal regions. Black stars indicate statistically significant differences between the control and heat-selection lines (GLMM: ***P<0.001). The data underlying this figure can be found at doi.org/10.5281/zenodo.22277897.

To investigate whether inversions facilitate or impede introgression and to account for the distinct evolutionary histories of the X chromosome and autosomes, we divided genomic regions into colinear (non-inverted) and inverted regions for both autosomes and the X chromosome. Autosomal colinear regions showed a significantly greater introgression in the heat-selection lines compared to the control lines (GLMM: z_1,10_ = 4.11, P < 0.001; Fig. 3, 4A, S2; Table S2). Autosomal inverted regions showed some introgression in three of the heat-selection replicate lines, while none of the control replicate lines exhibited any introgression (Fig. 3, 4A, S2; Table S2). However, this difference was not statistically significant (GLMM: z_1,10_ = 1.60, P = 0.110; Fig. 3, 4A, S2; Table S2). In contrast, the entire X chromosome, which contains three overlapping inversions, exhibited essentially no introgression from *D. montana* in either the heat-selection or control lines (GLMM: X_COL_: z_1,10_ = 1.24, P = 0.215; X_INV_: z_1,10_ = 0.12, P = 0.907; Fig. 3, 4A, S2; Table S2), suggesting that the X chromosome is largely impermeable to introgression. Furthermore, the variability in the mean HI was significantly higher among the heat-selection lines compared to the control lines in autosomal inverted regions (GLMM: z_1,10_ = 14.86, P < 0.001), while no significant differences were observed in other genome partitions (GLMM: Auto_COL_: z_1,10_ = -0.77, P = 0.441; X_COL_: z_1,10_ = -0.53, P = 0.598; X_INV_: z_1,10_ = -0.603, P = 0.546; Fig. 3, 4B, S2; Table S2). These results suggest that introgression is most prevalent in colinear autosomal regions but also occurs in inverted autosomal regions in some of the heat-selection lines.

To test whether introgression was adaptive, we analyzed correlations, blind to control and heat-exposed selection lines, between the proportion of fertile males after heat exposure and the mean HI from different genome partitions. Intriguingly, the fertility proportion and the mean HI showed a significant positive correlation for autosomal colinear regions (Fig. 5A), but not for other genomic regions (Fig. 5B-D). These results suggest that male fertility under heat stress improves with higher levels of introgression from the more heat-tolerant *D. montana* in autosomal colinear regions.

**Figure 5.**
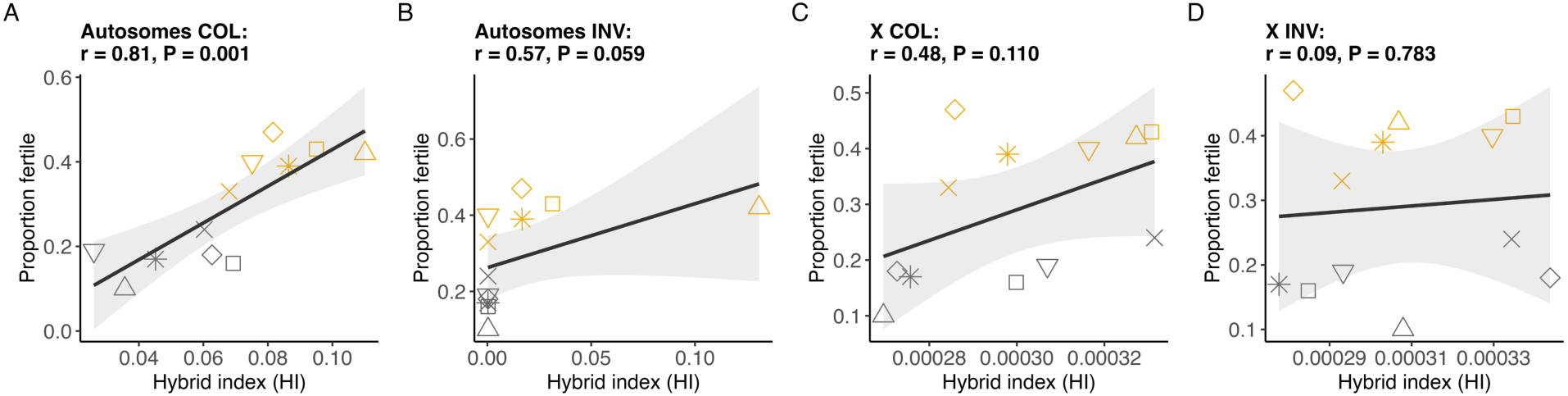
Correlation analysis between the proportion of fertile males after heat exposure and hybrid index (HI) of (A) autosomal colinear, (B) autosomal inverted, (C) X chromosomal colinear and (D) X chromosomal inverted regions. The data underlying this figure can be found at doi.org/10.5281/zenodo.22277897.

Finally, we identified genetic regions with an HI of 1, indicating that these regions were introgressed from the more heat-tolerant *D. montana* and fixed. In the control lines, none of SNPs were fixed, whereas the heat-selection lines exhibited seven fixed SNPs, ranging from 0 to 3 per line (mean = 1.17; Fig. 3B). None of these SNPs were shared across the six heat-selection replicate lines. Of the seven fixed SNPs, four were located within or near genes (Table S3). Some of these genes may have become fixed due to strong selection, and future work could examine the role of these genes in adaptation to heat stress.

### Heat stress selection can partly overcome long-standing barriers to gene flow

Given that the initial hybrid index at the start of the HER experiment was 25%, the amount of introgression from *D. montana* after a further three generations was relatively low in both the heat- selection (5.6%) and control (2.9%) lines, indicating the presence of effective barriers to gene flow. These barriers were identified using two complementary approaches. First, genetic regions with an HI of 0 − indicating no introgression from *D. montana* in any of the six control or six heat- selection replicate lines − indicate strong contemporary barriers to gene flow. Comparing these regions between the two sets of lines determines whether strong heat selection and the benefits of hybridization can overcome the strong barriers. Second, by analyzing the genomes of *D. flavomontana* and *D. montana* collected from natural populations using the demographically explicit genome scan gIMble, we identified candidate long-term barriers to gene flow, indicated by reduced long-term effective migration rates (*m_e_*) across genomic windows.

Approximately 95% of the SNPs exhibited no introgression (HI=0) in both inverted and colinear regions across both control and heat-selection lines on the X chromosome (Fig. 4C, Fig. 3, Table S2). Similarly, approximately 95% of the SNPs showed no introgression within the inverted autosomal regions in the control lines (Fig. 4C, Fig. 3, Table S2). However, this percentage decreased to 13% in heat-selection lines (Fig. 4C, Fig. 3, Table S2). In colinear autosomal regions, the proportion of SNPs with no introgression was 33% in the control lines and only 2% in the heat- selection lines (Fig. 4C, Fig. 3, Table S2). These results indicate that the X chromosome, with its complex rearrangements, acts as a strong barrier to introgression between the species. In contrast, in autosomal regions, strong heat stress selection overcame 95% of the barriers in colinear regions and 86% in inverted regions, suggesting that adaptive introgression occurs more easily in colinear regions.

The best-fit global demographic model for *D. flavomontana* and *D. montana* suggests that the species diverged ∼2.1 mya, with very low levels of unidirectional post-divergence gene flow from *D. montana* into *D. flavomontana* (0.04 migrants per generation; Table S4), consistent with previous findings (∼2.5 mya and 0.01 migrants per generation; [50]). Candidate long-term barrier windows were defined as positive Δ_B0_ values with a 5% false positive rate (FPR), where a history of reduced *m_e_* fits better than a model assuming the global *m_e_*. Overall, 36.8% (3,130 out of 8,499) of the windows were potential barriers to gene flow (Fig. 6A). By merging the overlapping windows, barriers were identified in 55.0% (78.2Mb out of 142.3Mb) of the genome, distributed across 304 regions (Fig. 6A). Notably, 85.2% (21.4 Mb out of 25.1 Mb) on autosomal inverted regions and 73.8% (11.8 Mb out of 16.0 Mb) on the X chromosomal inverted regions constituted candidate long-term barriers. In contrast, only 46.6% (41.2 Mb out of 88.3 Mb) of the autosomal colinear regions and 29.7% (3.9 Mb out of 13.0 Mb) of the X chromosomal colinear regions acted as barriers. The barriers were significantly more prevalent in inverted compared to colinear regions, both on autosomes (Chi-Square: X² = 88.62, P < 0.0001) and on the X chromosome (X² = 36.74, P < 0.0001).

**Figure 6.**
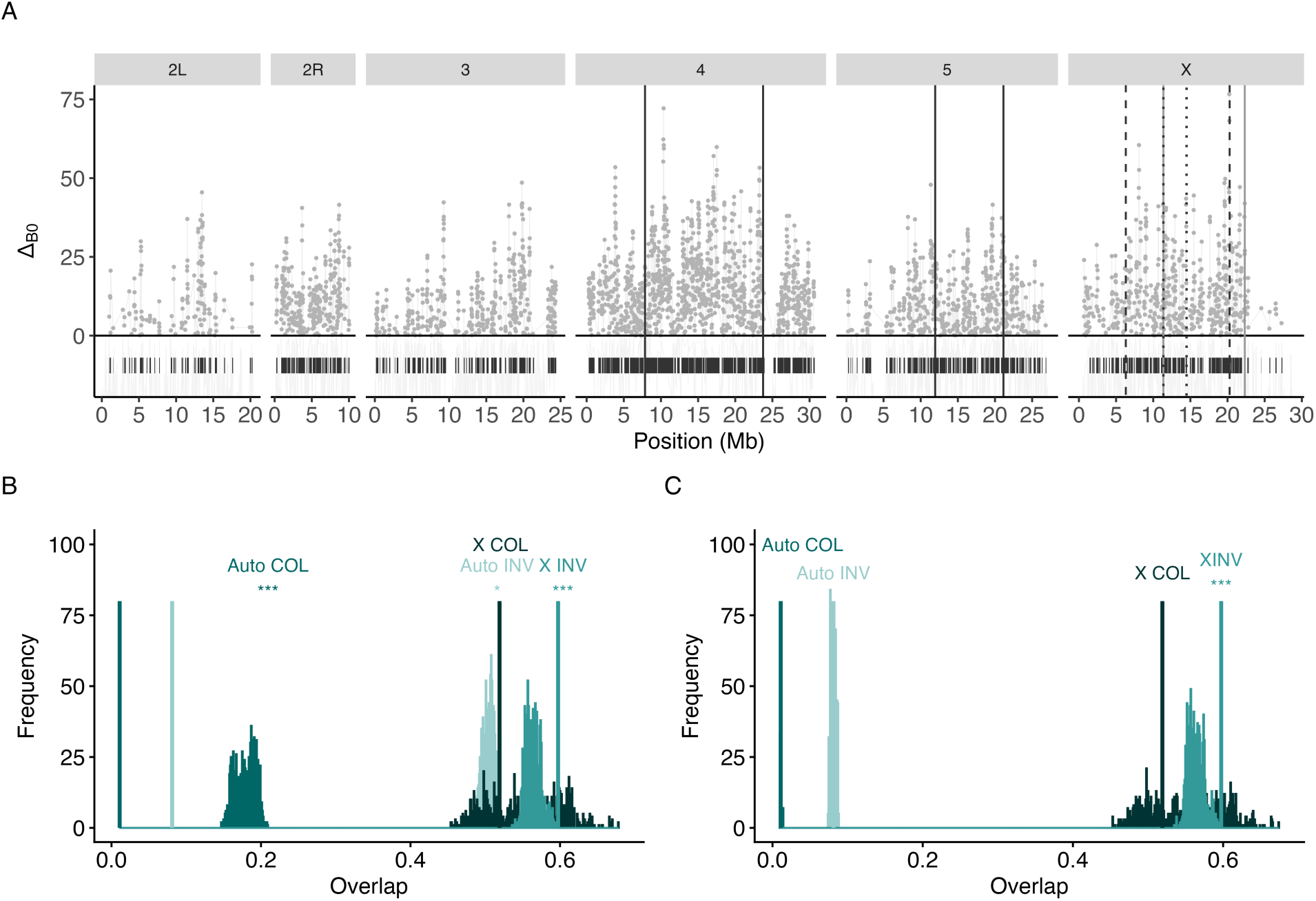
(A) Long-term barriers to gene flow between natural populations of *D. flavomontana* and *D. montana*. Dots indicate windows with Δ_B0_>0 and a false positive rate at 5%, i.e. candidate barriers to gene flow, where the history of reduced migration rate (*m_e_*) between the species fits better than a model assuming the global estimate. Barrier regions (overlapping windows with Δ_B0_>0 and FPR≤0.05) are marked with dark segments below 0. Vertical solid and dashed lines denote the breakpoints of the alternatively fixed chromosomal inversions between *D. flavomontana* and *D. montana*. The observed (vertical lines) and simulated (bootstrap distributions) overlap between long-term barriers and strong contemporary barriers (=SNPs showing no introgression; HI=0) in (B) the control and (C) the heat-selection treatment in colinear (COL) and inverted (INV) autosomal (Auto) and X chromosomal regions. If the observed value exceeds the simulated distribution, the strong contemporary barriers and long-term barriers overlap more than expected by chance (circular bootstrap: *P<0.05, **P<0.01, ***P<0.001). The data underlying this figure can be found at doi.org/10.5281/zenodo.22277897.

Strong contemporary barriers (SNPs showing no introgression, HI=0) observed in the control lines showed a significantly higher overlap with long-term barriers than expected by chance in all chromosome partitions (Circular bootstrap: Auto_COL_: P < 0.001; Auto_INV_: P = 0.024; X_INV_: P < 0.001), except in the colinear X chromosome (P = 0.685; Fig. 6B). In contrast, strong contemporary barriers observed in the heat-selection lines exhibited a significantly higher overlap with the long- term barriers only within the X chromosomal inversions (P < 0.001), and not in other regions (Auto_COL_: P = 0.867; Auto_INV_: P = 0.367; X_COL_: P = 0.686; Fig. 6C). These results suggest that at least some of the contemporary barriers observed in the lab crosses are of old origin, given that they overlap with the long-term barriers estimated to have started accumulating around the time when the species began to diverge, approximately 2.1 million years ago, and have contributed to maintaining species differentiation. Despite this, strong heat stress selection can partly outweigh the costs of any maladaptive incompatible loci and favor introgression, except within the X chromosomal inversions.

## Discussion

Adaptive introgression is considered an important source of genetic variation [2], particularly in the context of global warming [54]. However, only a few studies have experimentally tested the potential of adaptive introgression in response to warming temperatures [12,13], or examined the interplay between adaptive introgression, barrier loci, and recombination-suppressing chromosomal inversions. We focused on heat-induced male sterility given its link to current species distributions and increased extinction risk compared to survival traits [15,18]. We investigated these topics by conducting a hybridize, evolve, and re-sequence (HER) experiment by crossing the heat-sensitive *D. flavomontana* females with the more heat-tolerant *D. montana* males, backcrossing the F_1_ females with *D. flavomontana* males, and exposing heat-selection lines to transient heat for three generations, while maintaining control lines. Our findings suggest that strong heat stress selection can mitigate genetic barriers, promoting the transfer of heat-adapted alleles between species and enhancing heat tolerance. Introgression was most prevalent in colinear autosomal regions but also occurred to some extent in inverted autosomal regions. However, the X chromosome, with three overlapping inversions, resisted introgression due to persistent long-standing barriers, thereby maintaining species barriers in this part of the genome.

Parratt et al. [15] projected that suitable habitats for *D. flavomontana* will significantly shrink by 2080 due to climate change, as increasing temperatures cause heat-induced sterility in males, leading to population declines. Our findings suggest that introgression can facilitate the transfer of heat-adapted alleles from the more heat-tolerant *D. montana* into the heat-sensitive *D. flavomontana*, thereby enhancing heat tolerance despite persistent genetic barriers. The heat- selection lines exhibited approximately twice the levels of introgression and male fertility under heat stress compared to the control lines, and 1.5 times greater male fertility than the heat-sensitive parental *D. flavomontana*. Introgression primarily occurred in autosomal colinear regions, which showed a significant correlation with thermal fertility phenotypes across replicate lines. This suggests that higher levels of introgression from *D. montana* correspond to better male fertility under heat stress, indicating that introgression can provide a rapid source of adaptive genetic material under stressful conditions, helping to mitigate climate-induced population extinctions. While the thermal fertility phenotypes of the heat-selection lines did not match those of *D. montana*, suggesting limitations due to linked maladaptive loci or an insufficient selection period or population size, the introgression of some heat-adapted alleles would provide valuable additional time for new beneficial mutations to emerge and be selected. Although introgression between *D. flavomontana* and *D. montana* has occurred in nature (see Results and [50,55]), including the potential transfer of heat shock proteins [55], it is likely historical rather than ongoing [50]. Successful evolutionary rescue via adaptive introgression in the wild would require overcoming prezygotic barriers between the two species [51]. In the future, hybridization rates between them may increase if rising temperatures cause heat-induced sterility in conspecific males, which could lead females to be less choosy and accept heterospecific males to avoid the high costs of mate searching or the risk of remaining unmated [56,57]. Other studies have demonstrated the potential for adaptive introgression to improve viability under heat stress in an intraspecific context, with weaker barriers to gene flow [13,14]. Whether *D. flavomontana* populations can evolve resistance to heat-induced male sterility from standing genetic variation alone remains an open question, but introgression from *D. montana* offers a complementary source of genetic material, given that *D. montana* has roughly twice the genetic diversity and effective population size of *D. flavomontana* [50], and its males retain fertility at higher temperatures [15]. Extensive studies of natural systems would be needed to assess the relative importance of standing genetic variation and interspecific gene flow. The study also opens the door to other important questions for future research. While experimental studies suggest that introgression can improve stress resistance, what is the long-term fate of the introgressed material [6], and is it maladaptive for other traits, such as reducing fecundity or viability, or in other environmental contexts? Does introgression increase the risk of species fusion or extinction due to shared ecological niches or collapsed reproductive barriers [58,59], leading to a loss of biodiversity?

Chromosomal inversions suppress recombination across the inverted sequence [38,39], potentially facilitating the accumulation and maintenance of both adaptive loci [42] and barriers to gene flow [40,41]. Given that adaptive loci can become linked to maladaptive loci through reduced recombination within inversions, the role of the inversions in introgression is difficult to predict. As expected, we find that inversions strongly reduce recombination, not only within the inverted regions but also beyond the inversion breakpoints [60,61]. This is indicated by long blocks of similar hybrid indices throughout and beyond the inverted regions (Fig. 3). However, we do see some variability inside the inversions (e.g. inside chromosome 4 inversion, see Fig. 3B), presumably due to double crossovers [62] or gene conversion events [63]. In the control lines without heat stress selection, *D. montana* alleles from inverted regions were quickly eliminated across replicate lines. The evolution in these lines was predictable and consistent with patterns observed in previous studies of introgression in other species [27,33,64]. In contrast, we observed some introgression across autosomal inversions in some of the heat-selection lines, supporting the idea that these inversions may harbor adaptive alleles and that entire inverted regions can be transferred as “adaptive cassettes” from one species to another when the benefits of introgression outweigh the negative effects [46–49]. However, the levels of introgression within these inversions varied widely across the heat-selection replicate lines, making general predictions about the role of inversions in adaptive introgression challenging due to the randomness of recombination events. If an inversion can “survive” the first hybrid generations, some recombination within inversions and selection can purge linked incompatibilities over time, allowing adaptive alleles to escape from any linked deleterious alleles.

Although the heat-selection lines exhibited higher levels of introgression compared to the control lines (5.6% vs. 2.9%), introgression was relatively low in both overall, indicating the presence of substantial barriers to gene flow. We identified contemporary barriers in lab crosses and compared these with long-term barriers inferred using demographic modelling and made three key observations. First, there were fewer barriers in colinear autosomal regions compared to inverted ones, and under heat stress, these barriers were more effectively overcome in colinear regions (95%) than in inverted regions (86%). These findings suggest that the benefits of introgression under heat stress can outweigh its negative effects in both colinear and inverted autosomal regions, but this effect is more pronounced in colinear regions. Second, the X chromosome with three overlapping inversions was entirely sheltered from introgression in both the heat-selection and control lines. This is likely due to several factors: i) sex chromosomes are often disproportionately involved in the build-up of barriers to gene flow, particularly BDMIs, due to factors such as faster-X evolution and meiotic drive [29,65–67], ii) the fixation probability of inversions is higher on the X chromosome compared to autosomes [68,69], and iii) complex rearrangements reduce recombination more effectively than simpler rearrangements [70]. Overall, sex-linked chromosomal inversions may play a crucial role in maintaining species integrity in the face of gene flow, supporting previous findings in *D. flavomontana* and *D. montana* [34] and in other species [44]. Finally, we found a significant overlap between contemporary barriers observed in the control lines and long-term barriers in both inverted and colinear genomic regions. A similar overlap was observed between contemporary barriers in natural hybrid populations and long-term barriers inferred from demographic modelling in *Iphiclides* butterflies [29] and *Formica* red wood ants [30]. However, this overlap was less strong in the heat-selection lines, suggesting that strong heat stress selection can mitigate even long-standing genetic barriers. Overall, identifying the same genetic barriers using different approaches provides strong evidence for accurate barrier mapping.

Our design of the HER experiment and the study system can present some challenges in interpreting the results. Given the low number of hybrid offspring emerging after heat exposure, we used nearly all available males to establish the next generation (see Materials and methods). This could have led to an unequal number of fertile males in the heat-selection and control lines, potentially resulting in a lower effective population size and higher levels of genetic drift in the heat-selection lines. However, it is unlikely that random effects would consistently increase both the amount of introgression and improve fitness in the heat-selection lines compared to the control lines. Drift would instead generate inconsistent patterns of introgression. Furthermore, while selection on hybrids could result from purifying selection acting on low-fitness alleles from lab-adapted strains, this explanation would predict increased fitness and introgression in both the heat- selection and control lines. We observe increased fitness and introgression only in the heat- selection lines, indicating that response to selection is due to the experimental heat conditions rather than general lab adaptation. An alternative explanation for improved fertility of the heat- selection lines could be existing genetic variation in *D. flavomontana* or transgenerational plasticity. However, the low genetic diversity of isofemale strains, the better heat tolerance of heat- selection lines compared to *D. flavomontana*, the generally weak thermal plasticity in insects [71], and the correlation of improved thermal fertility limits with the extent of introgression from *D. montana* in colinear regions suggest that adaptive introgression was the primary mechanism. Finally, although we observed increased introgression from *D. montana* in the heat-selection lines compared to the control lines, pinpointing actual targets of selection is challenging during the early stages of introgression due to the hitchhiking of neutral SNPs in large linkage blocks [72], and because polygenic adaptation typically involves minor shifts in allele frequencies [73]. While a higher number of hybrid generations may reduce the overall amount of introgression and diminish the statistical power to compare treatments, they could also allow recombination to separate adaptive loci from barrier and neutral loci, potentially enabling the identification of exact causal regions for adaptation.

In conclusion, experimental studies enable direct tests of hybridization, adaptive introgression and factors maintaining species’ integrity. Our study provides evidence for a successful evolutionary rescue via adaptive introgression in response to climate change in a relatively old species pair, which we hope will encourage similar studies in other systems and address some of the open questions.

## Material and methods

### Hybridize-evolve-and-re-sequence (HER) experiment to detect adaptive introgression and contemporary barriers to gene flow

#### Isofemale strains used in the experiment

In the HER experiment, we used two *D. flavomontana* and two *D. montana* isofemale strains, which were established from fertilized females collected in sympatry in Jackson, Wyoming, USA (43°26’N, 110°50’W) in 2013. The isofemale strain codes are flaJX13F37, flaJX13F38, monJX13F3, and monJX13F48. Since their establishment, these strains were maintained under laboratory conditions at 20±1 °C in continuous light to prevent females from entering reproductive diapause (∼75 generations). Emerging flies were sexed under light CO_2_ anesthesia within 72 hours. Flies were transferred into fresh malt vials once a week and used in experiments at 18-22 days old, when they are sexually mature.

#### Heat tolerance experiments prior to the HER experiment

Prior to the HER experiment, we identified the temperatures that sterilize 50-80% of *D. flavomontana* males but have a weaker effect on *D. montana* males. Based on previous research in these species [15], we selected six temperatures for testing: 20 °C (control), 28 °C, 29 °C, 30 °C, 31 °C, and 32 °C. Groups of males were exposed to these temperatures for four hours in a Julabo circulator bath and then returned to 20±1 °C, following the protocol by Parratt et al. [15]. Each male was then placed in a malt vial with two sexually mature, virgin females for seven days, after which all flies were discarded. Males were scored as “fertile” if larvae were present in the vials. Each temperature was tested 2-4 times, except 20 °C which was tested once, with each replicate including 11-39 males. Based on the results (see Results), 31 °C and 32 °C induce 50- 80% sterility in *D. flavomontana* males. Consequently, we tested the sterilizing effects of these temperatures and the control temperature of 20 °C on *D. montana* males. Both 31 °C and 32 °C were tested twice, and 20 °C was tested once, with each replicate consisting of 11-35 *D. montana* males per strain. Additionally, we examined the effect of heat on female fertility by exposing females of the *D. flavomontana* and *D. montana* strains to the same temperatures as the males for four hours, and placing each heated female with two sexually mature, virgin males for seven days to score fertility. Each temperature was tested once, with each replicate consisting of 9-24 females per strain.

We tested i) whether fertility of *D. flavomontana* males decreased with increasing temperatures and ii) whether *D. montana* males had greater fertility after heat exposure at 31 °C and 32 °C compared to *D. flavomontana* males. We used a generalized linear model (GLM) with a Binomial distribution, having fertile and sterile males as the response variable, and temperature or species as the explanatory variables. Statistical analyses were conducted using *glm* function in base R (v4.4.1) and Rstudio (v2024.04.2).

#### The HER experiment

To start the HER experiment, we crossed 50 *D. flavomontana* females with 50 *D. montana* males for each of the two strain pairs (flaJX13F37xmonJX13F3 and flaJX13F38xmonJX13F48; Fig. 1). Because F_1_ hybrid males are sterile [34,51], we backcrossed F_1_ hybrid females to *D. flavomontana* males. Specifically, for each strain pair we set up five replicate backcross lines, each consisting of 20 randomly selected virgin F_1_ females mass-mated to 20 virgin *D. flavomontana* males (Fig. 1). From the resulting first backcross (BC_1_) generation, we randomly selected 60 virgin females and 60 virgin males per replicate line and split them evenly between the heat-selection and control treatments (30 of each sex per replicate per treatment), following Griffiths et al. [13]. Starting with these BC_1_ females and males, we conducted three generations of selection. In each generation, virgin males and females in the heat-selection treatment were separately exposed to heat for four hours: 31 °C in the first generation, 31.5 °C in the second, and 32 °C in the third. In the control treatment, flies were not exposed to heat but were otherwise handled identically. The 30 virgin females and 30 virgin males per replicate line, either heat-exposed or non-heat-exposed depending on the treatment, were placed together in a malt vial to establish the next generation. These species have a generation time of approximately seven weeks. Figure 1 outlines the HER experiment.

Note that we could not test the fertility of individual males of the heat-selection lines after heat exposure in each generation and select only fertile males for the next generation due to two major issues: i) there were simply not enough males for picking specific males; instead nearly all emerging males were exposed to heat and used to establish the next generation, and ii) if a male was found to be sterile, this could have been due to either heat-induced sterility or genetic incompatibilities, with the latter possibly depending on the specific male-female pair. Therefore, it was important to allow the flies to mate freely rather than assigning pairs. This study design may lead to an unequal number of males in the heat-selection and control lines, potentially resulting in a lower effective population size and a higher effect of genetic drift in the heat-selection lines. However, replicating the heat-selection and control lines helps in estimating the effect of heat- selection in hybrids. Additionally, recombination occurs only in *Drosophila* females, meaning that the recombination events should be roughly the same under both heat-selection and control treatments.

To reduce the effects of acclimation on the results, the flies used for phenotypic tests and for DNA extractions and sequencing were collected from the fourth generation of intercrossed BC_1_ hybrids (BC_1_F_4_), i.e. the unexposed offspring of the heat-exposed BC_1_F_3_ generation (see Fig. 1). During the three generations of heat selection, one of the selection replicates of flaJX13F37xmonJX13F3 strain pair and two of the selection replicates of flaJX13F38xmonJX13F48 strain pair went extinct. Given that all control lines performed well, these extinctions likely resulted from heat stress rather than genetic incompatibilities. Consequently, due to the loss of some heat-selection replicates, the maximum number of lines in a balanced design was three heat-selection and three control lines per strain pair (Fig. 1), totalling six heat-selection and six control lines altogether. Further details of these procedures are provided in the following sections.

#### Heat tolerance experiments of the heat-selection and control lines after the HER experiment

The fourth generation of intercrossed BC_1_ males (BC_1_F_4_ ♂) of the heat-selection and control lines were phenotyped to test whether heat selection had improved males’ ability to tolerate heat (measured as heat-induced sterility). Males were exposed to 32 °C for four hours in a Julabo circulator bath, and then returned to 20±1 °C. Each male was placed in a malt vial with two sexually mature, virgin BC_1_F_4_ females of the respective heat-selection and control lines for seven days, after which all flies were discarded. Males were scored as “fertile” if larvae were present in the vials. The experiment was replicated 2-4 times, the number of males per replicate and line ranging from 8 to 45 (mean 22, median 21), depending on the number of flies available.

We tested differences in male fertility using a GLMM with a Binomial distribution, having fertile and sterile males as the response variable, the treatment (heat-selection, control) as the explanatory variable, and the strain pair (flaJX13F37×monJX13F3, flaJX13F38×monJX13F48) as the random effect. Statistical analyses were conducted using the *glmmTMB* function from glmmTMB R package [74].

#### Sequencing of the heat-selection and control lines after the HER experiment

We collected 20 fourth generation of intercrossed BC_1_ females (BC_1_F_4_♀) from each of the six heat-selection and six replicate lines, as well as from the four parental isofemale strains (flaJX13F37, flaJX13F38, monJX13F3, monJX13F48). DNA extractions were performed on pools of 10 flies using cetyltrimethylammonium bromide (CTAB) solution with RNAse treatment, followed by Phenol-Chloroform-Isoamyl alcohol (25:24:1) and Chloroform-Isoamyl alcohol (24:1) washing steps, and ethanol precipitation. The genomic DNA was quantified using a Qubit fluorometer (Thermo Fisher Scientific). An equal amount of DNA (1600ng) from two pools of 10 females was combined to create the final sample, which included DNA from 20 females per pooled sample. These steps were conducted at the University of St Andrews (Scotland, UK) in 2024. The quality-checked DNA was then used to generate libraries, and 150bp paired-end (PE) reads were sequenced on an Illumina NovaSeq X Plus Series at Novogene, aiming for 40X coverage (each of the 20 females in a pool has two chromosome copies, resulting in 40 chromosome copies). The observed mean coverage for the heat-selection and control lines and for the parental strains ranged from 31X to 44X. Further details about the Illumina raw reads are given in Table S5.

#### Mapping and variant calling of the heat-selection and control lines

Pooled Illumina PE reads were trimmed for adapter contamination and low-quality bases using fastp v0.21.0 [75] and then mapped to the *D. flavomontana* reference genome [50] using BWA mem v0.7.17 with read group information [76]. All analyses were performed using the *D. flavomontana* reference genome because previous comparative analyses of the *D. flavomontana* and *D. montana* genomes indicated that reference bias has a minimal impact on the results [34,50]. The alignments were sorted with SAMtools v1.10 [77] and PCR duplicates were marked with sambamba v0.7.0 [78]. BAM-files were filtered for mapping quality of >20 using SAMtools. Allele counts for each sample at each genomic position were obtained with SAMtools mpileup, using options to retain reads with a mapping quality of >20 and sites with a base quality of >20. The resulting mpileup file was used for variant calling against the *D. flavomontana* reference genome using the heuristic SNP calling software, PoolSNP [79]. In PoolSNP, we specified a minimum count of 5 to call a SNP and a minimum coverage of 8. For a maximum coverage, we considered positions within the 95% coverage percentile for a given sample and chromosome. Variant calling detected a total of 3,897,771 biallelic SNPs.

#### Analysis of hybrid index (HI) of the heat-selection and control lines

The expected amount of genetic material transferred from females of one species into the other, i.e. hybrid index (HI), is 50% in F_1_ hybrids and 25% after one round of backcrossing. The first round of heat selection was performed on the first backcross (BC_1_) generation hybrids, which have 25% *D. montana* ancestry. After three rounds of heat selection, changes in introgression can be measured using the HI and compared between the heat-selection and control treatments.

We computed the HI based on species-diagnostic SNPs, which are variants differentially fixed between the parental *D. flavomontana* and *D. montana*, following Poikela et al. [34]. To obtain a reliable dataset, we defined differentially fixed SNPs using both pool-sequenced data from two strains of each species and individually sequenced wild-caught *D. flavomontana* and *D. montana* females (the results remained consistent even when defining the differentially fixed SNPs using only the pool-sequenced data from the two strains of each species). Details about the wild-caught samples are provided in Table S6 and the “Identifying long-term barriers to gene flow in natural populations of the species” section. In total, 700,751 diagnostic SNPs were identified. For each diagnostic SNP, allele frequencies of the *D. montana* alleles were calculated in each of the six heat- selection and six control lines. The mean HI and the absolute deviation from the mean HI were then computed and analysed for the control and heat-selection lines and for the colinear and inverted autosomal and X chromosomal regions. Inversions and the X chromosome were analysed separately due to their distinct evolutionary histories [34,50,80]. The mean HI between the heat- selection and control treatments was analysed using a GLMM with a beta distribution, with the mean HI as the response variable, the treatment (heat-selection, control) as the explanatory variable, and the strain pair (flaJX13F37×monJX13F3, flaJX13F38×monJX13F48) as the random effect. The absolute deviation was analyzed similarly, using either a Gamma or Gaussian distribution, depending on the normality of the data. Statistical analyses were conducted using the *glmmTMB* function from glmmTMB R package [74]. Finally, to test if introgression was potentially adaptive, we examined the correlations between the mean HI and the mean fertility proportions across the control and heat-selection replicate lines. These correlations were tested separately for each chromosome partition using either the Pearson or Spearman correlation test, depending on the normality of the data.

Additionally, we identified two further categories of SNPs: i) SNPs with an HI of 1 in each line, indicating that these regions were introgressed from the more heat-tolerant *D. montana* and fixed, and ii) SNPs showing no introgression from *D. montana* (HI=0) in any of the six control and six heat-selection lines, indicating strong contemporary barriers. Given that the selection lasted only for three generations, we do not expect many SNPs to be fixed for the *D. montana* allele (HI=1), nor do we necessarily expect these fixed SNPs to be shared across replicate lines. Finally, we extracted genes carrying SNPs fixed for the *D. montana* allele (HI=1), either within the gene body or in the 2000bp flanking regions, and blasted them against *Drosophila melanogaster* and *Drosophila virilis* RefSeq proteins (BLASTp v2.15.0+; Table S3; [81]). We chose *D. melanogaster* for its superior annotation, and *D. virilis* for its closer relation to *D. flavomontana* and *D. montana*.

### Locations and presence of chromosomal inversions

Previous comparative analysis of contiguous genome assemblies, along with PacBio long-read and Illumina short-read data, revealed that *D. flavomontana* and *D. montana* differ by five large chromosomal inversions [50]. Three of these inversions overlap and are located on the X chromosome, and one each on autosomes 4 and 5 (inversion breakpoints are shown in Table S1). The presence of these inversions was visually confirmed by examining the orientation, insert size, and clipped reads of paired-end Illumina pools mapped to the *D. flavomontana* reference genome around each breakpoint using the Interactive Genomics Viewer (IGV, [82]). As anticipated, the *D. montana* reads showed reversed orientation, extended insert size, and clipped reads around the breakpoints, whereas the *D. flavomontana* reads did not. Due to the complex overlap of the three inversions on the X chromosome, the read orientation around some X inversion breakpoints was not reversed, but insert size was extended. Since chromosome 4 of the *D. flavomontana* reference genome was scaffolded, the presence of the inversion on chromosome 4 was verified by mapping the parental pools to the *Drosophila lummei* reference genome, as described in Poikela et al. [50]. IGV plots are presented in Fig. S3.

### Identifying long-term barriers to gene flow in natural populations of the species

#### Genomic samples of wild-caught females

*D. flavomontana* samples were collected from Jackson, Wyoming, USA (43°26’N, 110°50’W), and stored in 70% EtOH at -20 °C in 2013. DNA extractions were performed for eight individual females, each the offspring of a different wild-caught female, using cetyltrimethylammonium bromide (CTAB) solution with RNAse treatment, Phenol-Chloroform-Isoamyl alcohol (25:24:1) and Chloroform-Isoamyl alcohol (24:1) washing steps and ethanol precipitation at the University of St Andrews (Scotland, UK) in 2023. Quality-checked DNA was then used to generate libraries, and 150bp paired-end (PE) reads were sequenced on an Illumina NovaSeq 6000 at Novogene. Additionally, sequences from one individually sequenced *D. flavomontana* female and three *D. montana* females were obtained from GenBank under Bioprojects PRJNA828433 and PRJNA939085. These samples were collected from the same location and time as the *D. flavomontana* samples described above. The mean coverage for the *D. flavomontana* samples sequenced in this study ranged from 12X to 35X, while the previously sequenced *D. flavomontana* and *D. montana* samples had a coverage range of 57X to 140X. Further details about the samples and Illumina raw reads are provided in Table S6. Note that the two *D. flavomontana* and *D. montana* isofemale strains used in the HER experiment were derived from two of the wild-caught *D. flavomontana* females and two wild-caught *D. montana* females sequenced here.

#### Demographic modelling using gIMble

The individually sequenced wild-caught *D. flavomontana* and *D. montana* females were used to identify regions of reduced gene flow, i.e. candidate barrier regions, across the genome using gIMble (genome-wide IM blockwise likelihood estimation toolkit; [36]). Briefly, gIMble detects barriers to gene flow through heterogeneity in migration rate (*m_e_*), while accounting for background selection via heterogeneity in effective population sizes (*N_e_*). For this analysis, Illumina PE reads were trimmed for adapter contamination and low-quality bases using fastp v0.21.0 [75]. The trimmed reads were then mapped to the *D. flavomontana* reference genome [50] using BWA mem v0.7.17 with read group information [76]. The alignments were sorted with SAMtools v1.10 [77] and PCR duplicates were marked with sambamba v0.7.0 [78]. The resulting BAM files were used for variant calling with freebayes v1.3.6 [83], resulting in a total of 7,186,955 biallelic SNPs. The variants were filtered using gIMble “preprocess” module.

Using gIMble, we have recently demonstrated that the best-fit demographic model for *D. flavomontana* and *D. montana* is an isolation with migration (IM) model with post-divergence gene flow occurring from *D. montana* to *D. flavomontana* (forward in time) [50]. To confirm this phylogeographic history, we compared three demographic scenarios: strict divergence without post-divergence gene flow (DIV, *m_e_* = 0), and isolation with migration (IM) models with both gene flow directions between the species (IM_monàfla_, IM_flaàmon_, forward in time; see Table S4). This initial model comparison focused on non-repetitive intergenic sequences and introns shorter than 80bp to minimize the direct effects of selection (see [55]). Regions with fixed inversions and the X chromosome were excluded from this analysis due to their different evolutionary histories compared to other genomic regions [50]. Colinear autosomal regions, ending at inversion breakpoints (Table S1), were combined across the genome, as these regions are expected to share the same evolutionary history. Data were summarized using block-wise site frequency spectrum (bSFS) with a block length of 64b with k_max_ values of 2. bSFS-based analyses assume no recombination within blocks and a constant mutation rate (μ) across them. We used μ = 2.8×10−9 per site per generation, based on an estimate of the spontaneous mutation rate in *Drosophila melanogaster* [84]. The estimates of divergence time (*T*) were converted into absolute time using t = *T* × 2*N_e_* × g, where *N_e_* = θ/(4μ) and g is generation time, assuming one generation per year [53,85].

To identify long-term barriers to gene flow, we used the best-fit model in the “local” mode of gIMble to identify regions of reduced *m_e_* across the genome. Following Laetsch et al. [36], 500 blocks of 64bp each were grouped into windows, resulting in a nominal span of 32 kb. Since only non-coding, non-repetitive blocks were used, the actual window span was greater than 32 kb (mean window span 75.5kb). We explored variation in *N_e_*s and *m_e_* across the windows by searching for parameter combinations on a 12 x 12 x 12 x 20 grid (*N_e Dfla_*, *N_e Dmon_*, *N_e_* _ancestral_, *m_e_*). The estimate of *T* was obtained from the global analysis and fixed here, as it is assumed to be a global event shared across the genome. Local support for reduced *m_e_* (window-wise variation in *m_e_* shown in Fig. S4) was measured via positive Δ_B0_ [36]. The false positive rate (FPR) at 5% of Δ_B0_ was estimated using msprime by simulating 100 window-wise datasets with globally fixed *m_e_*, while accounting for variation in *N_e_* by assuming the best composite log-likelihood *N_e_* parameter inferred for each window. Only windows where Δ_B0_ in the real data exceeded the largest value in the Δ_B0_ distribution from the simulated data were labeled as barriers. In the simulations, a constant recombination rate was used, based on one crossover per chromosome during female meiosis (recombination occurs only in females), resulting in a map length of 25cM per chromosome. Finally, the significant overlapping windows with reduced gene flow (Δ_B0_>0 and FPR≤0.05) were merged to obtain barrier regions. The number of non-overlapping barrier and non-barrier windows was compared between the inverted and colinear regions on autosomes and on the X chromosome using a Chi-Square test to determine whether barriers are enriched within the inversions.

### Overlap analysis between the contemporary barriers observed in lab crosses and the long-term barriers identified using gIMble

To test whether SNPs showing no introgression (HI=0) across the heat-selection or control lines − indicative of the strongest contemporary barriers to gene flow observed in the laboratory crosses − were associated with long-term barriers identified using gIMble (proportion of HI=0 SNPs within gIMble barrier and non-barrier windows), we employed a circular resampling approach from Ebdon et al. [29] (https://github.com/LohseLab/circular_bootstrap/tree/main). This approach involves circularizing each chromosome and sampling with random offsets, generating null distributions for any genomic measure, while maintaining the same distribution and spacing of genomic windows as the barrier windows inferred by gIMble. It detects whether observed SNPs showing no introgression are significantly enriched in long-term barrier windows compared to non-barrier windows. The observed estimates were compared to the distributions of means of 1000 resampled datasets, following Ebdon et al. [29]. This overlap analysis was conducted separately for colinear and inverted autosomal and X chromosomal regions, and for both heat-selection and control treatments.

## Acknowledgements

This research was supported by a grant from Jenny and Antti Wihuri Foundation to NP, and MGR is funded by NERC UK (NE/V001566/1). We would like to thank Prof. Anneli Hoikkala and Dr. Maaria Kankare who kindly provided the wild-caught *D. flavomontana* samples and *D. flavomontana* isofemale strains for this project. We also thank the Finnish Center for Scientific Computing (CSC) for computing resources used in this project.

## Author contributions

All authors contributed to the conceptualization and methodology of the research. NP performed DNA extractions, experiments, and data analyses. MGR supervised the research. NP drafted the first version of the manuscript, and all authors reviewed and edited the draft. NP and MGR funded the study.

## Data availability

Raw reads from previously sequenced wild-caught *D. montana* and *D. flavomontana* females were obtained from the SRA under BioProjects PRJNA939085 and PRJNA828433. The reference genome was obtained from Dryad (https://doi.org/10.5061/dryad.4f4qrfjft). Newly sequenced raw reads of wild-caught *D. flavomontana*, as well as the parental and hybrid lines, are publicly available at the SRA under BioProject PRJNA1215028. All code and phenotypic data are publicly available on Zenodo (10.5281/zenodo.22277897).

## Ethics declaration

*D. flavomontana* and *D. montana* are not endangered, and the flies were collected along watersides on public lands outside National and State parks, where insect collecting does not require permits in the USA (The Wilderness Act of 1964, section 6302.15).

## Supporting information

**Table S1.** Genomic coordinates for the breakpoints of alternatively fixed chromosomal inversions between *D. flavomontana* and *D. montana*. The coordinates were obtained from Poikela et al. [50]

**Table S2.** Hybrid indices (HIs) and strong barriers to gene flow in the control and heat-selection treatments. The table shows information on the number of diagnostic SNPs (= alternatively fixed SNPs between the species), the mean and standard deviation of the hybrid indices (HIs), and the number and percentage of SNPs with no introgression from *D. montana* (HI = 0) across the control or heat-selection replicate lines.

**Table S3.** *Drosophila melanogaster* and *Drosophila virilis* orthologues of genes in the heat-selection lines that contain SNPs introgressed and fixed from the more heat-tolerant *D. montana* (HI=1), located within the gene body or in the 2000bp flanking regions. No SNPs were introgressed and fixed in the control lines.

**Table S4.** Demographic parameters estimated by gIMble. Parameters for effective population sizes (*N_e_*), divergence time (*T* in years/generations) and migration rate (*m_e_*) between *D. flavomontana* and *D. montana* estimated from 64bp blocks under the strict divergence (DIV) and isolation with migration (IM) models with both gene flow directions. *m_e_* estimates correspond to M (=4N_e_*m_e_*) individuals per generation (forward in time).

**Table S5.** Details of the sampling site, geographic coordinates and sampling year as well as sequencing information for *D. flavomontana* and *D. montana* isofemale strains used in the hybridize, evolve, and re- sequence (HER) experiment, and for the resulting heat-selection and control lines.

**Table S6.** Details of the sampling site, geographic coordinates and sampling year as well as sequencing information for *D. flavomontana* and *D. montana* samples used in demographic modelling. Four of the samples were first published by Poikela et al. [50] and Tahami et al. [86] and the rest in the current study.

**Figure S1.** Proportion of fertile *D. flavomontana* and *D. montana* females and males after heat exposure at 28-32 °C and at the control temperature of 20 °C. Flies were kept at each temperature for four hours, after which their fertility was assessed. The data underlying this figure can be found at 10.5281/zenodo.22277897.

**Figure S1.** Variation in hybrid index (HI) across the genome between the control and heat-selection treatments. (A) The relative difference in HI across the genome between the heat-selection and control lines, where positive values indicate greater introgression in the heat-selection compared to the control lines and negative values greater introgression in the control compared to the heat-selection lines. Vertical solid and dashed grey lines in all plots represent the breakpoints of the alternatively fixed chromosomal inversions between *D. flavomontana* and *D. montana*. (B) The difference in the mean HI between the respective heat- selection and control lines in colinear (COL) and inverted (IV) autosomal and X chromosomal regions. The data underlying this figure can be found at 10.5281/zenodo.22277897.

**Figure S1.** Inversion breakpoints of alternatively fixed inversions between *D. montana* and *D. flavomontana* pools. Illumina paired-end re-sequencing data of both species were mapped against *D. flavomontana* reference genome, and the data is illustrated with Integrative Genomics Viewer (IGV) [82]. Due to inversion differences between the species, *D. montana* reads are split, the insert size of its paired reads deviates from the expected (red reads), and paired reads are in reversed orientation (blue and turquoise reads). Note that not all X inversion breakpoints exhibit reversed orientation due to the complex overlap of these inversions (see Table S1). Since chromosome 4 of the *D. flavomontana* reference genome was scaffolded, the presence of the inversion on chromosome 4 was verified by mapping both species to the *Drosophila lummei* reference genome (breakpoints 1,568,294 and 15,689,984 in the *D. lummei* genome equal to 23,745,872 and 7,855,585 in the *D. flavomontana* genome, respectively; Table S1), as described in Poikela et al. [50]. The data underlying this figure can be found at 10.5281/zenodo.22277897.

**Figure S1.** Variation in migration rate (*m_e_*) across sliding windows between natural populations of *D. flavomontana* and *D. montana*. Vertical solid and dashed lines denote the breakpoints of the alternatively fixed chromosomal inversions between *D. flavomontana* and *D. montana*. The data underlying this figure can be found at 10.5281/zenodo.22277897.

