## Supplementary material for "Adaptive introgression between two *Drosophila* species enhances heat tolerance despite barriers to gene flow"

### Supplementary Tables

Table S1. Genomic coordinates for the breakpoints of alternatively fixed chromosomal inversions between *D. flavomontana* and *D. montana*. The coordinates were obtained from Poikela et al. [1]

| Scaffold | Scaffold length | Start | End | Size (bp) |
|---|---|---|---|---|
| X_chromosome | 28975156 | 6309450 | 20284519 | 13975070 |
| X_chromosome | 28975156 | 11367083 | 22321964 | 10954882 |
| X_chromosome | 28975156 | 11368055 | 14487633 | 3119579 |
| 4_chromosome | 30698533 | 7855585 | 23745872 | 15890288 |
| 5_chromosome | 27217941 | 11949884 | 21134310 | 9184427 |

Table S2. Hybrid indices (HIs) and strong barriers to gene flow in the control and heat-selection treatments. The table shows information on the number of diagnostic SNPs (= alternatively fixed SNPs between the species), the mean and standard deviation of the HIs, and the number and percentage of SNPs with no introgression from *D. montana* (HI = 0) across the control or heat-selection replicate lines.

| Genomic region | Diagnostic SNPs | Control lines: Mean HI ± SD | Heat-selection lines: Mean HI ± SD | SNPs with no introgression across control lines (HI=0) | SNPs with no introgression across heat-selection lines (HI=0) |
|---|---|---|---|---|---|
| All | 700,751 | 2.88 ± 0.98 | 5.58 ± 1.75 | 414,560 (59.2%) | 181,469 (25.9%) |
| Autosomes colinear | 403,688 | 4.98 ± 1.70 | 8.60 ± 1.50 | 131,255 (32.5%) | 6,462 (1.6%) |
| Autosomes inverted | 131,562 | 0.03 ± 0.002 | 3.27 ± 4.95 | 125,473 (95.4%) | 17,290 (13.1%) |
| X colinear | 72,650 | 0.03 ± 0.002 | 0.03 ± 0.002 | 69,368 (95.5%) | 69,288 (95.4%) |
| X inverted | 92,851 | 0.03 ± 0.003 | 0.03 ± 0.002 | 88,464 (95.3%) | 88,429 (95.2%) |

Table S3. *Drosophila melanogaster* and *Drosophila virilis* orthologues of genes in the heat-selection lines that contain SNPs introgressed and fixed from the more heat-tolerant *D. montana* (HI=1), located within the gene body or in the 2000bp flanking regions. No SNPs were introgressed and fixed in the control lines.

| Chromosome | SNP position | Gene ID | *D. melanogaster* orthologue | REFSEQ ID | E-value | *D. virilis* orthologue | REFSEQ ID | E-value |
|---|---|---|---|---|---|---|---|---|
| 2L | 8445419 | - | - | - | - | - | - | - |
| 2L | 8445477 | - | - | - | - | - | - | - |
| 3 | 10612878 | g6867.t1 | uncharacterized protein Dmel_CG32137 | NP_729936.1 | 0.0 | bicaudal D-related protein homolog | XP_070064005.1 | 0.0 |
| 3 | 10985433 | g6917.t1 | Mob2 | NP_729716.1 | 0.0 | MOB kinase activator-like 2 | XP_032290013.1 | 0.0 |
| 3 | 11523112 | g6959.t1 | glucuronyltransferase P | NP_648448.1 | 0.0 | galactosylgalactosylxylosylprotein 3-beta-glucuronosyltransferase P | XP_002048547.2 | 0.0 |
| 4 | 28932682 | - | - | - | - | - | - | - |
| 5 | 15161128 | g12423.t1 | uncharacterized protein Dmel_CG43341 | NP_001246190.1 | 1.46E-31 | serine/threonine-protein kinase tousled-like 1 | XP_015030261.1 | 2.32E-72 |

Table S4. Demographic parameters estimated by gIMble. Parameters for effective population sizes ($N_e$), divergence time (*T* in years/generations) and migration rate ($m_e$) between *D. flavomontana* and *D. montana* estimated from 64bp blocks under the strict divergence (DIV) and isolation with migration (IM) models with both gene flow directions. $m_e$ estimates correspond to M (=$4N_e m_e$) individuals per generation (forward in time).

| Model | Ancestral Ne | *D. fla* A Ne | *D. mon* B Ne | *T* (years) | *m* (M) | ΔlnCL |
|---|---|---|---|---|---|---|
| DIV | 1291727 | 349841 | 671803 | 2016061 | - | 9737 |
| IM *D. mon --> D. fla* | 1244138 | 338270 | 684919 | 2145402 | 1.28E-08 (0.04) | 0 |
| IM *D. fla --> D. mon* | 1260477 | 352864 | 665369 | 2122941 | 1.04E-08 (0.01) | 4098 |

Table S5. Details of the sampling site, geographic coordinates and sampling year as well as sequencing information for *D. flavomontana* and *D. montana* isofemale strains used in the hybridize, evolve, and re-sequence (HER) experiment, and for the resulting heat-selection and control lines.

| Illumina 150bp paired-end reads<br>Species | Population | Collection coordinates and year | Strain ID | # lanes | # reads (M) | Bases (Gb) | Mean coverage (observed) | Library | Technique | quencing facil | Sequencing year |
|---|---|---|---|---|---|---|---|---|---|---|---|
| *D. flavomontana* | Jackson | 43˚26'N; 110˚50'W; 1 857 m | flaJX13F37 | 2 | 60.0 | 9.0 | 35.2 | Truseq | NovaSeq X Plus Series | Novogene | 2024 |
| | Wyoming, USA | 2013 | flaJX13F38 | 1 | 55.5 | 8.3 | 32.9 | Truseq | NovaSeq X Plus Series | Novogene | 2024 |
| *D. montana* | Jackson | 43˚26'N; 110˚50'W; 1 857 m | monJX13F3 | 2 | 63.3 | 9.5 | 38.9 | Truseq | NovaSeq X Plus Series | Novogene | 2024 |
| | Wyoming, USA | 2013 | monJX13F48 | 1 | 59.2 | 8.9 | 35.6 | Truseq | NovaSeq X Plus Series | Novogene | 2024 |
| Heat-selection line | Lab crosses | | flaJX13F37xmonJX13F3: Sel2 | 1 | 67.6 | 10.1 | 40.7 | Truseq | NovaSeq X Plus Series | Novogene | 2024 |
| (hybrids) | | | flaJX13F37xmonJX13F3: Sel3 | 1 | 65.4 | 9.8 | 39.6 | Truseq | NovaSeq X Plus Series | Novogene | 2024 |
| | | | flaJX13F37xmonJX13F3: Sel4 | 1 | 61.7 | 9.3 | 37.3 | Truseq | NovaSeq X Plus Series | Novogene | 2024 |
| | | | flaJX13F38xmonJX13F48: Sel1 | 1 | 55.0 | 8.3 | 31.6 | Truseq | NovaSeq X Plus Series | Novogene | 2024 |
| | | | flaJX13F38xmonJX13F48: Sel3 | 1 | 52.6 | 7.9 | 31.0 | Truseq | NovaSeq X Plus Series | Novogene | 2024 |
| | | | flaJX13F38xmonJX13F48: Sel4 | 1 | 57.3 | 8.6 | 34.3 | Truseq | NovaSeq X Plus Series | Novogene | 2024 |
| Control line | Lab crosses | | flaJX13F37xmonJX13F3: Cont2 | 2 | 61.3 | 9.2 | 35.6 | Truseq | NovaSeq X Plus Series | Novogene | 2024 |
| (hybrids) | | | flaJX13F37xmonJX13F3: Cont3 | 1 | 54.9 | 8.2 | 34.4 | Truseq | NovaSeq X Plus Series | Novogene | 2024 |
| | | | flaJX13F37xmonJX13F3: Cont4 | 1 | 56.9 | 8.5 | 32.6 | Truseq | NovaSeq X Plus Series | Novogene | 2024 |
| | | | flaJX13F38xmonJX13F48: Cont1 | 1 | 57.7 | 8.7 | 34.4 | Truseq | NovaSeq X Plus Series | Novogene | 2024 |
| | | | flaJX13F38xmonJX13F48: Cont3 | 1 | 73.7 | 11.1 | 43.9 | Truseq | NovaSeq X Plus Series | Novogene | 2024 |
| | | | flaJX13F38xmonJX13F48: Cont4 | 1 | 56.7 | 8.5 | 33.7 | Truseq | NovaSeq X Plus Series | Novogene | 2024 |

Table S6. Details of the sampling site, geographic coordinates and sampling year as well as sequencing information for *D. flavomontana* and *D. montana* samples used in demographic modelling. Four of the samples were first published by Poikela et al. [1] and Tahami et al. [2] and the rest in the current study.

| Illumina 150bp paired-end reads<br>Species | Population | Collection coordinates | Collection year | Strain ID | # lanes | # reads (M) | Bases (Gb) | Mean coverage | Library | Technique | Sequencing facility | Sequencing year | Data first published |
|---|---|---|---|---|---|---|---|---|---|---|---|---|---|
| *D. flavomontana* | Jackson | 43˚26'N; 110˚50'W | 2013 | flaJX13F10 | 1 | 14.2 | 2.1 | 11.8 | Truseq | NovaSeq 6000 | Novogene | 2023 | current study |
| | Wyoming, USA | Altitude 1 857 m | 2013 | flaJX13F11 | 1 | 26.4 | 4.0 | 22.0 | Truseq | NovaSeq 6000 | Novogene | 2023 | current study |
| | | | 2013 | flaJX13F12 | 1 | 24.1 | 3.6 | 20.1 | Truseq | NovaSeq 6000 | Novogene | 2023 | current study |
| | | | 2013 | flaJX13F15 | 1 | 29.1 | 4.4 | 24.2 | Truseq | NovaSeq 6000 | Novogene | 2023 | current study |
| | | | 2013 | flaJX13F19 | 1 | 29.3 | 4.4 | 24.4 | Truseq | NovaSeq 6000 | Novogene | 2023 | current study |
| | | | 2013 | flaJX13F31 | 1 | 16.2 | 2.4 | 13.5 | Truseq | NovaSeq 6000 | Novogene | 2023 | current study |
| | | | 2013 | flaJX13F37 | 1 | 42.3 | 6.3 | 35.2 | Nextera | HiSeq 4000 | Edinburgh Genomics | 2017 | Poikela et al. 2024 |
| | | | 2013 | flaJX13F38 | 2 | 168.1 | 25.2 | 140.1 | Truseq | NovaSeq 6000 | Novogene | 2023 | current study |
| | | | 2013 | flaJX13F8 | 1 | 27.1 | 4.1 | 22.6 | Truseq | NovaSeq 6000 | Novogene | 2023 | current study |
| *D. montana* | Jackson | 43˚26'N; 110˚50'W | 2013 | monJX13F3 | 2 | 130.0 | 19.5 | 108.3 | Nextera | HiSeq 4000 | Edinburgh Genomics | 2017 | Poikela et al. 2024 |
| | Wyoming, USA | Altitude 1 857 m | 2013 | monJX13F41 | 1 | 79.6 | 11.9 | 66.3 | Truseq | HiSeq X-Ten | Novogene | 2020 | Tahami et al. 2024 |
| | | | 2013 | monJX13F48 | 1 | 68.0 | 10.2 | 56.6 | Truseq | HiSeq X-Ten | Novogene | 2020 | Tahami et al. 2024 |

#### Supplementary Figures

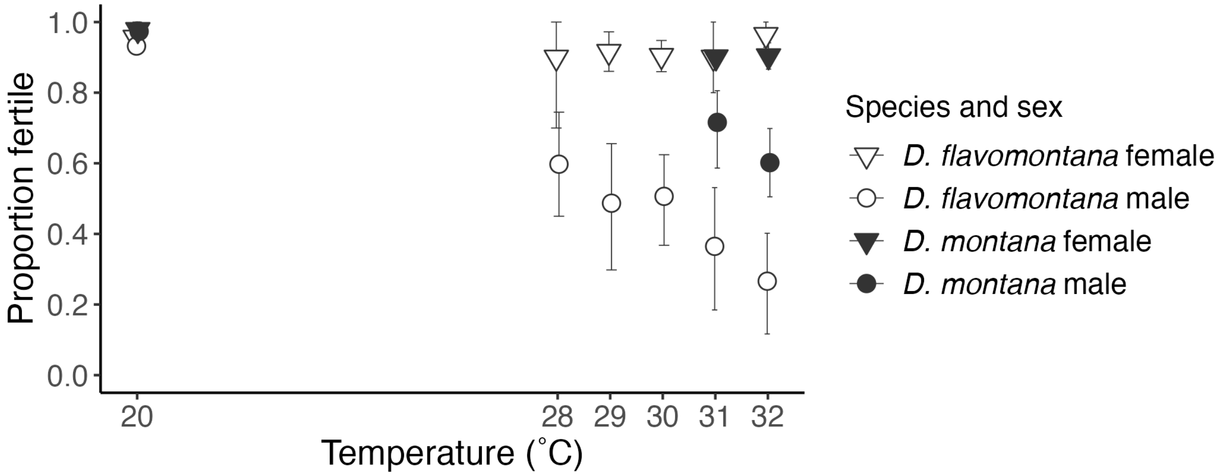


Figure S1. Proportion of fertile *D. flavomontana* and *D. montana* females and males after heat exposure at 28-32 °C and at the control temperature of 20 °C. Flies were kept at each temperature for four hours, after which their fertility was assessed. The data underlying this figure can be found at doi.org/10.5281/zenodo.22277897.

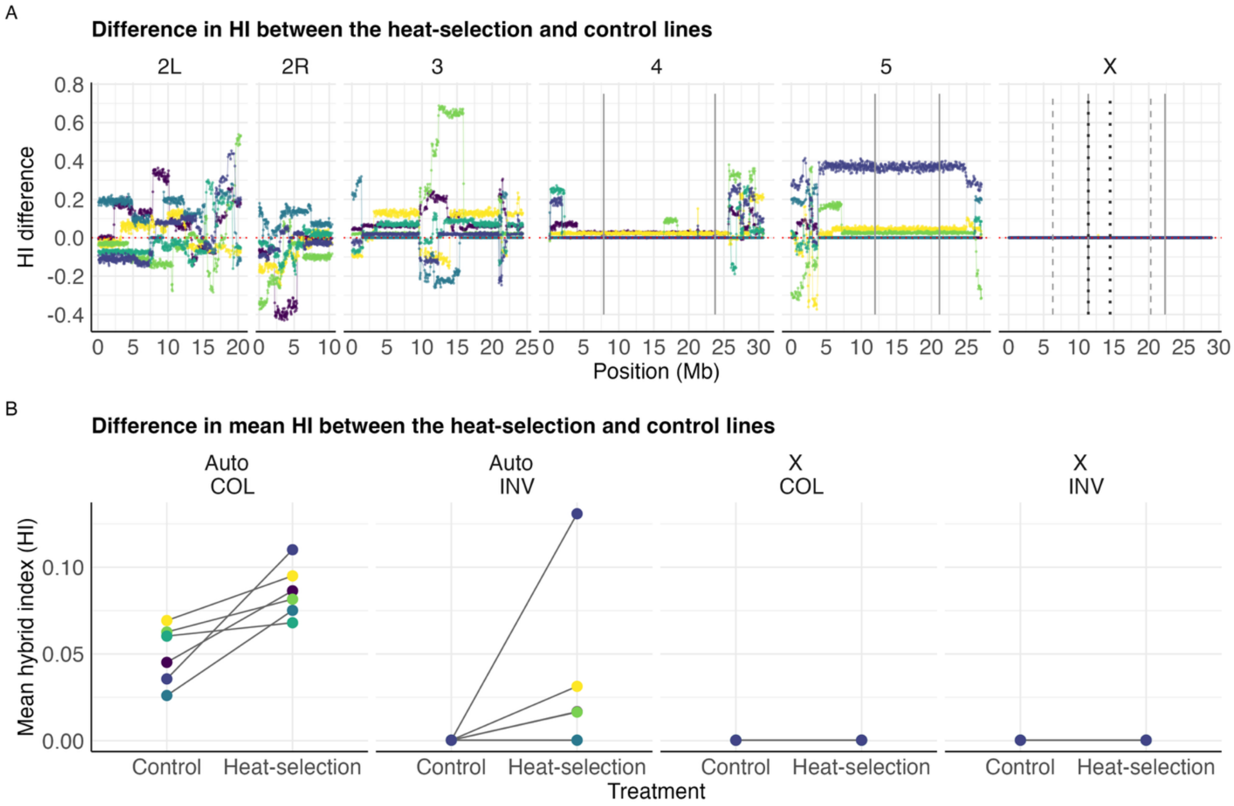


Figure S2. Variation in hybrid index (HI) across the genome between the control and heat-selection treatments. (A) The relative difference in HI across the genome between the heat-selection and control lines, where positive values indicate greater introgression in the heat-selection compared to the control lines and negative values greater introgression in the control compared to the heat-selection lines. Vertical solid and dashed grey lines in all plots represent the breakpoints of the alternatively fixed chromosomal inversions between *D. flavomontana* and *D. montana*. (B) The difference in the mean HI between the respective heat-selection and control lines in colinear (COL) and inverted (IV) autosomal and X chromosomal regions. The data underlying this figure can be found at doi.org/10.5281/zenodo.22277897.

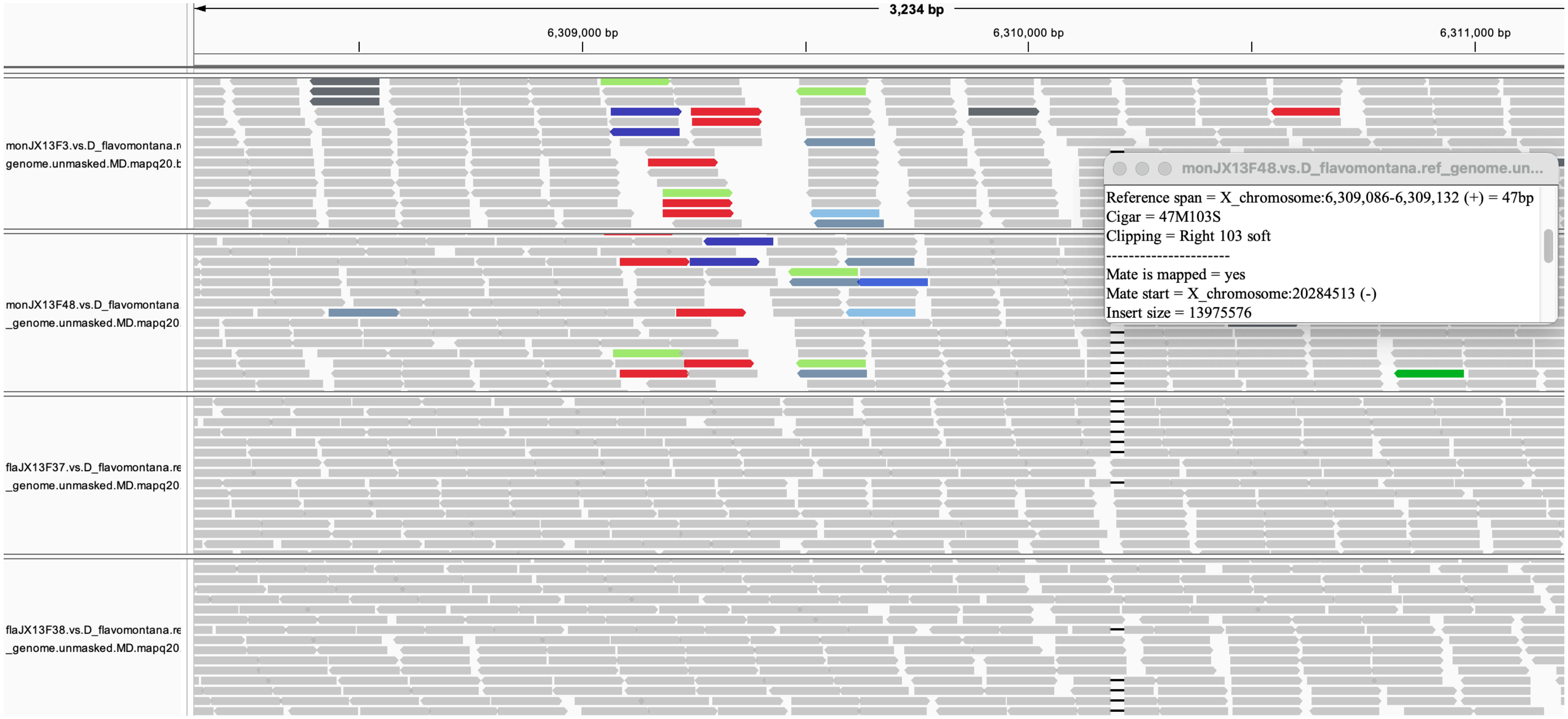
3,234 bp
6,309,000 bp
6,310,000 bp
6,311,000 bp
monJX13F3.vs.D_flavomontana.r
genome.unmasked.MD.mapq20.b
monJX13F48.vs.D_flavomontana
_genome.unmasked.MD.mapq20
flaJX13F37.vs.D_flavomontana.re
_genome.unmasked.MD.mapq20
flaJX13F38.vs.D_flavomontana.re
_genome.unmasked.MD.mapq20
monJX13F48.vs.D_flavomontana.ref_genome.un...
Reference span = X_chromosome:6,309,086-6,309,132 (+) = 47bp
Cigar = 47M103S
Clipping = Right 103 soft
----------------------
Mate is mapped = yes
Mate start = X_chromosome:20284513 (-)
Insert size = 13975576

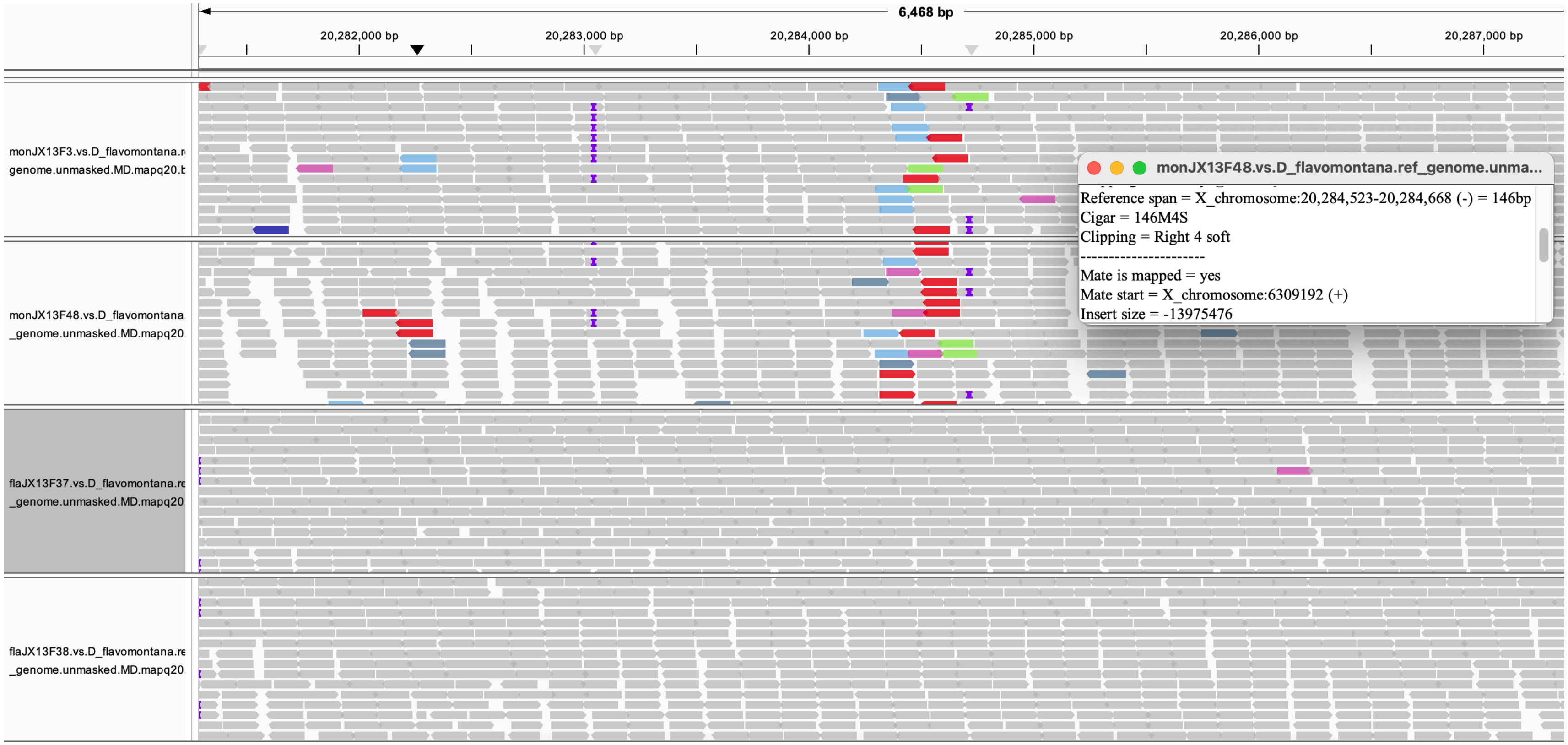
6,468 bp
20,282,000 bp
20,283,000 bp
20,284,000 bp
20,285,000 bp
20,286,000 bp
20,287,000 bp
monJX13F3.vs.D_flavomontana.r
genome.unmasked.MD.mapq20.b
monJX13F48.vs.D_flavomontana
_genome.unmasked.MD.mapq20
flaJX13F37.vs.D_flavomontana.re
_genome.unmasked.MD.mapq20
flaJX13F38.vs.D_flavomontana.re
_genome.unmasked.MD.mapq20
monJX13F48.vs.D_flavomontana.ref_genome.unma...
Reference span = X_chromosome:20,284,523-20,284,668 (-) = 146bp
Cigar = 146M4S
Clipping = Right 4 soft
Mate is mapped = yes
Mate start = X_chromosome:6309192 (+)
Insert size = -13975476

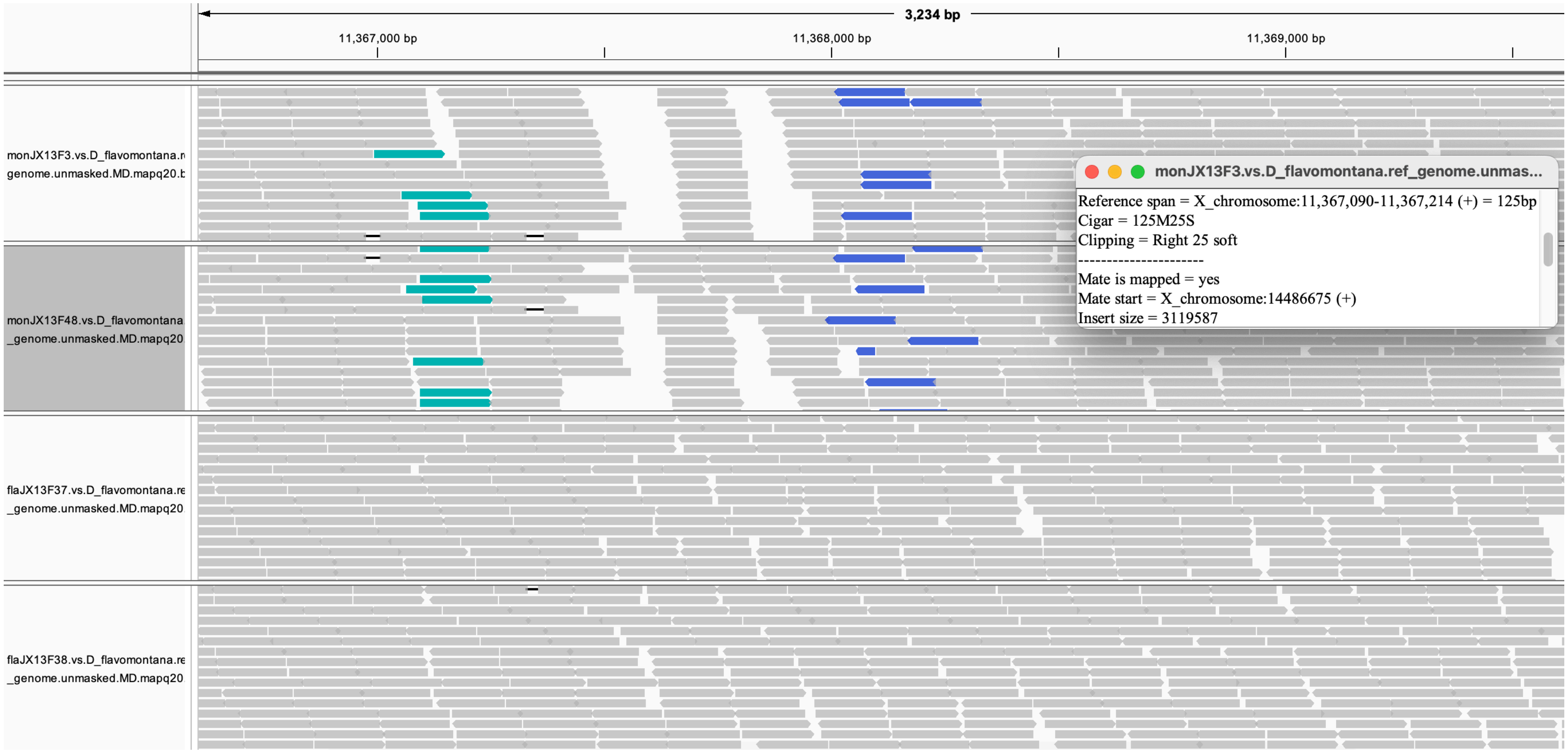
3,234 bp
11,367,000 bp
11,368,000 bp
11,369,000 bp
monJX13F3.vs.D_flavomontana.r
genome.unmasked.MD.mapq20.b
monJX13F48.vs.D_flavomontana
_genome.unmasked.MD.mapq20
flaJX13F37.vs.D_flavomontana.re
_genome.unmasked.MD.mapq20
flaJX13F38.vs.D_flavomontana.re
_genome.unmasked.MD.mapq20
monJX13F3.vs.D_flavomontana.ref_genome.unmas...
Reference span = X_chromosome:11,367,090-11,367,214 (+) = 125bp
Cigar = 125M25S
Clipping = Right 25 soft
Mate is mapped = yes
Mate start = X_chromosome:14486675 (+)
Insert size = 3119587

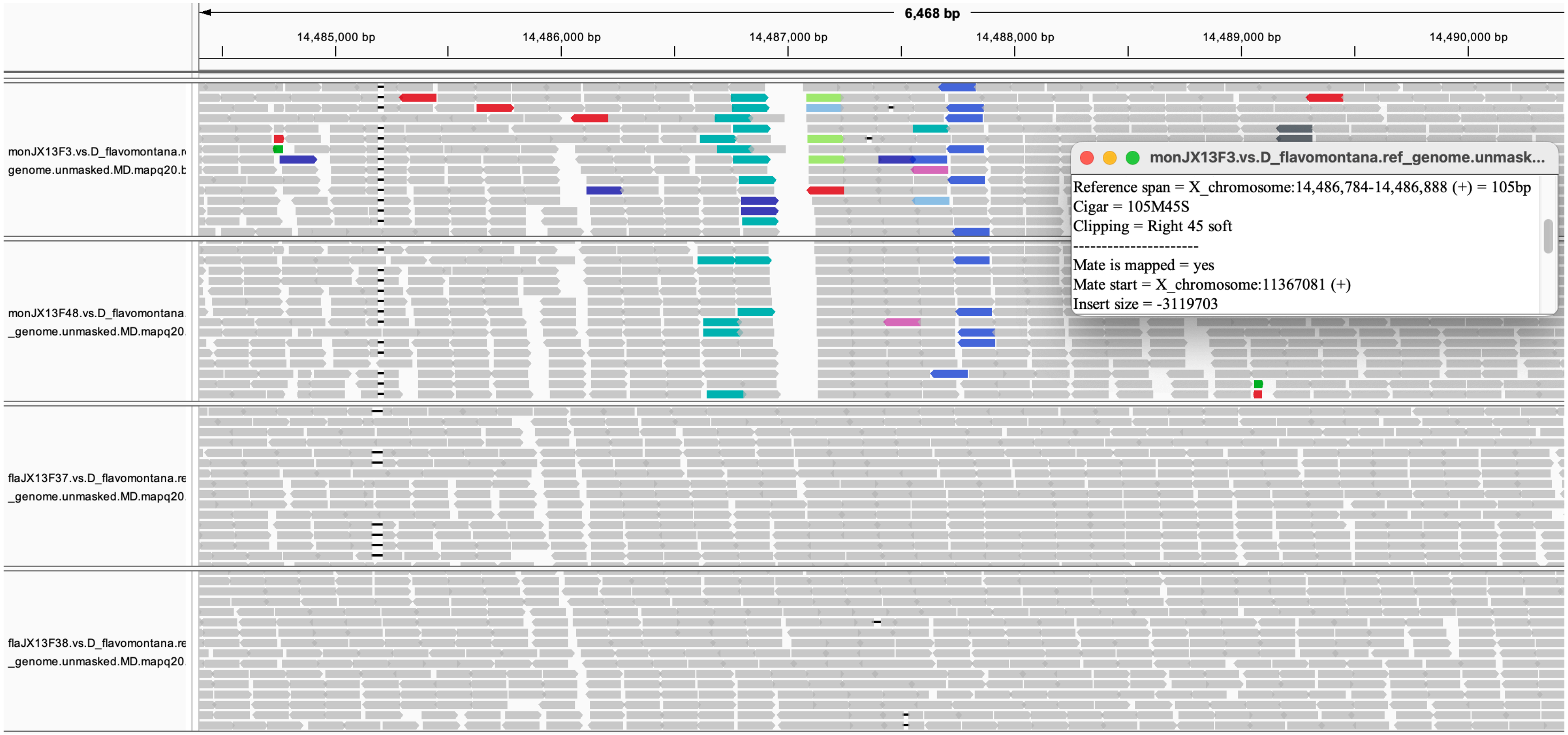
6,468 bp
14,485,000 bp
14,486,000 bp
14,487,000 bp
14,488,000 bp
14,489,000 bp
14,490,000 bp
monJX13F3.vs.D_flavomontana.r
genome.unmasked.MD.mapq20.b
monJX13F48.vs.D_flavomontana
_genome.unmasked.MD.mapq20
flaJX13F37.vs.D_flavomontana.re
_genome.unmasked.MD.mapq20
flaJX13F38.vs.D_flavomontana.re
_genome.unmasked.MD.mapq20
monJX13F3.vs.D_flavomontana.ref_genome.unmask...
Reference span = X_chromosome:14,486,784-14,486,888 (+) = 105bp
Cigar = 105M45S
Clipping = Right 45 soft
----------------------
Mate is mapped = yes
Mate start = X_chromosome:11367081 (+)
Insert size = -3119703

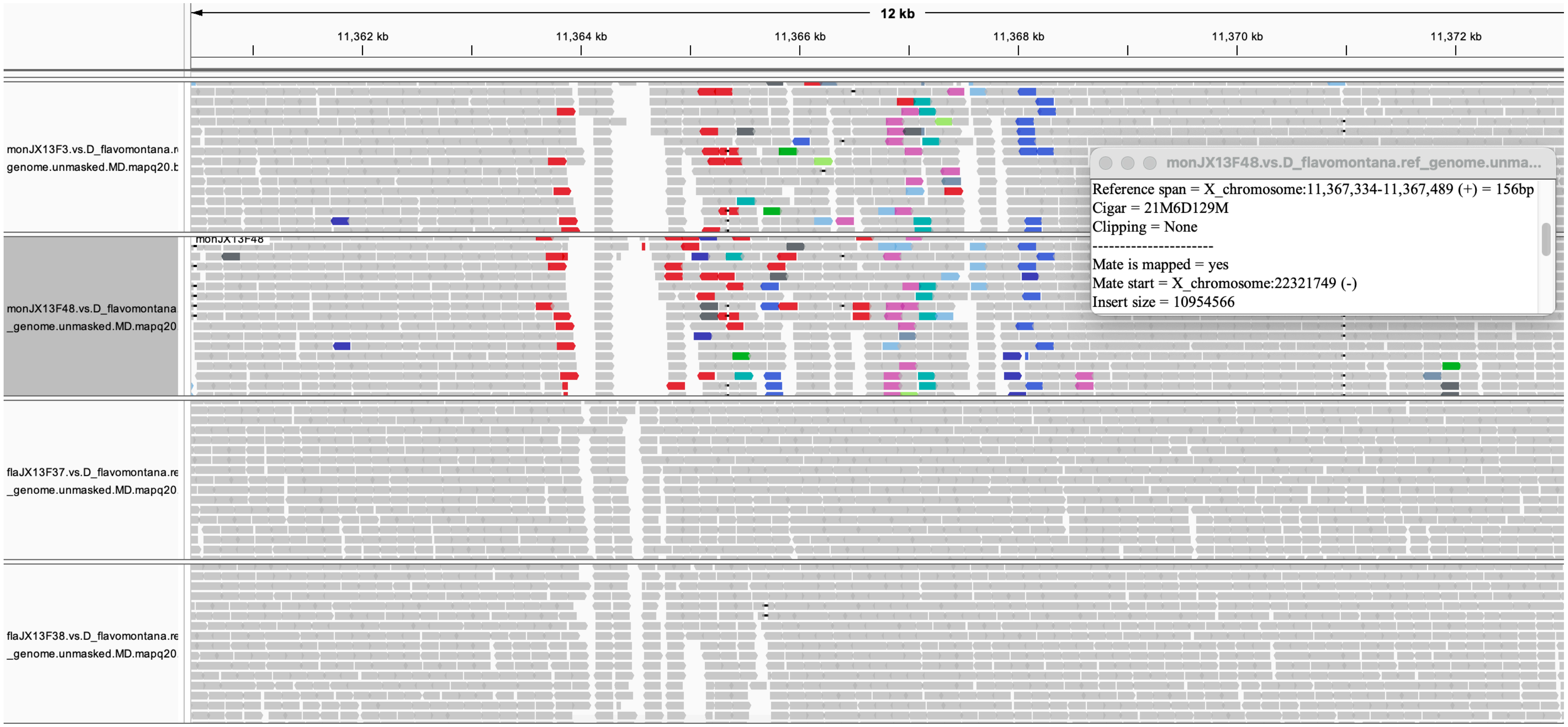
12 kb
11,362 kb
11,364 kb
11,366 kb
11,368 kb
11,370 kb
11,372 kb
monJX13F3.vs.D_flavomontana.r
genome.unmasked.MD.mapq20.b
monJX13F48.vs.D_flavomontana
_genome.unmasked.MD.mapq20
monJX13F48
flaJX13F37.vs.D_flavomontana.re
_genome.unmasked.MD.mapq20
flaJX13F38.vs.D_flavomontana.re
_genome.unmasked.MD.mapq20
monJX13F48.vs.D_flavomontana.ref_genome.unma...
Reference span = X_chromosome:11,367,334-11,367,489 (+) = 156bp
Cigar = 21M6D129M
Clipping = None
----------------------
Mate is mapped = yes
Mate start = X_chromosome:22321749 (-)
Insert size = 10954566

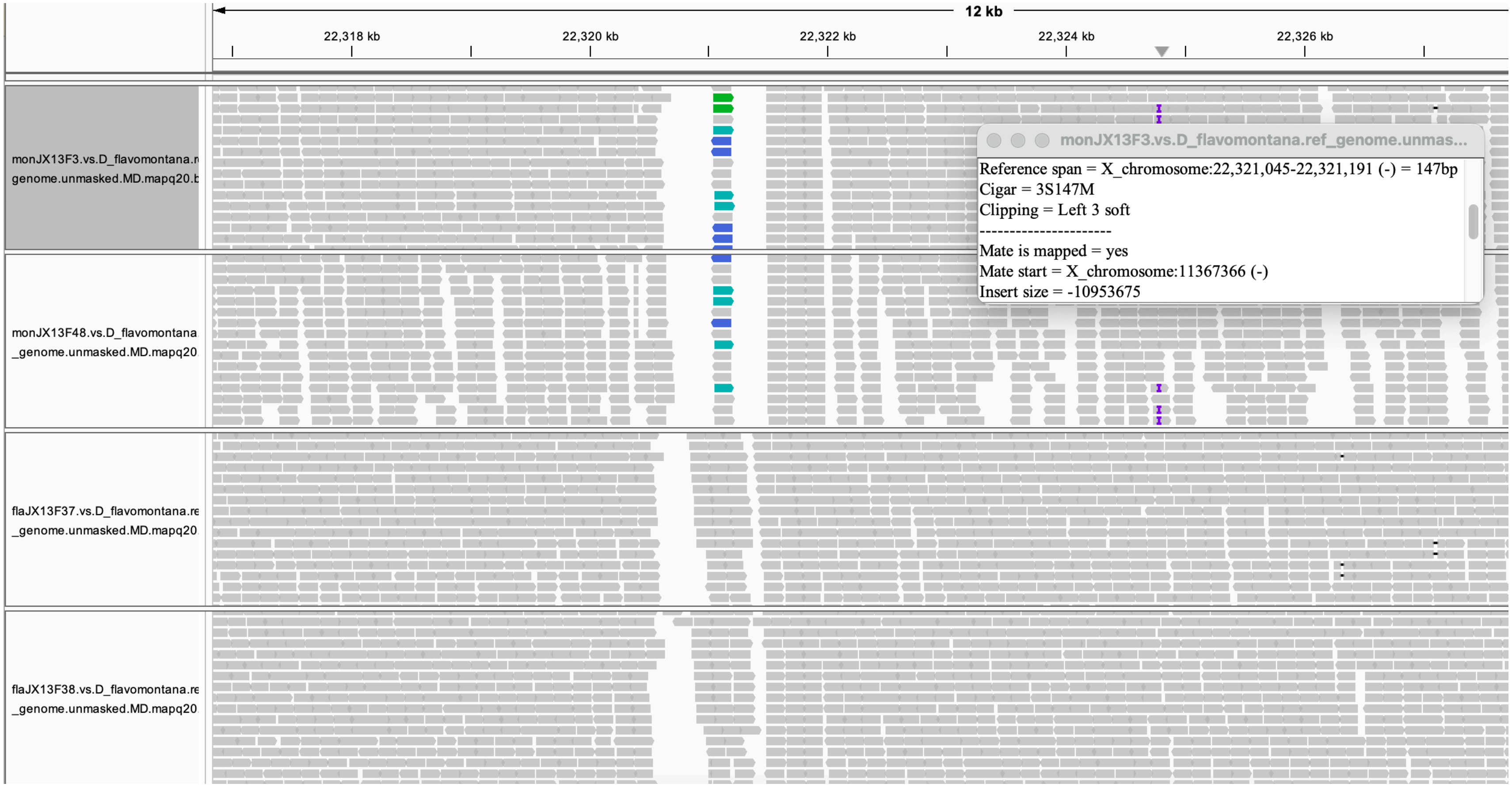
12 kb
22,318 kb
22,320 kb
22,322 kb
22,324 kb
22,326 kb
monJX13F3.vs.D_flavomontana.r
genome.unmasked.MD.mapq20.b
monJX13F48.vs.D_flavomontana
_genome.unmasked.MD.mapq20
flaJX13F37.vs.D_flavomontana.re
_genome.unmasked.MD.mapq20
flaJX13F38.vs.D_flavomontana.re
_genome.unmasked.MD.mapq20
monJX13F3.vs.D_flavomontana.ref_genome.unmas...
Reference span = X_chromosome:22,321,045-22,321,191 (-) = 147bp
Cigar = 3S147M
Clipping = Left 3 soft
----------------------
Mate is mapped = yes
Mate start = X_chromosome:11367366 (-)
Insert size = -10953675

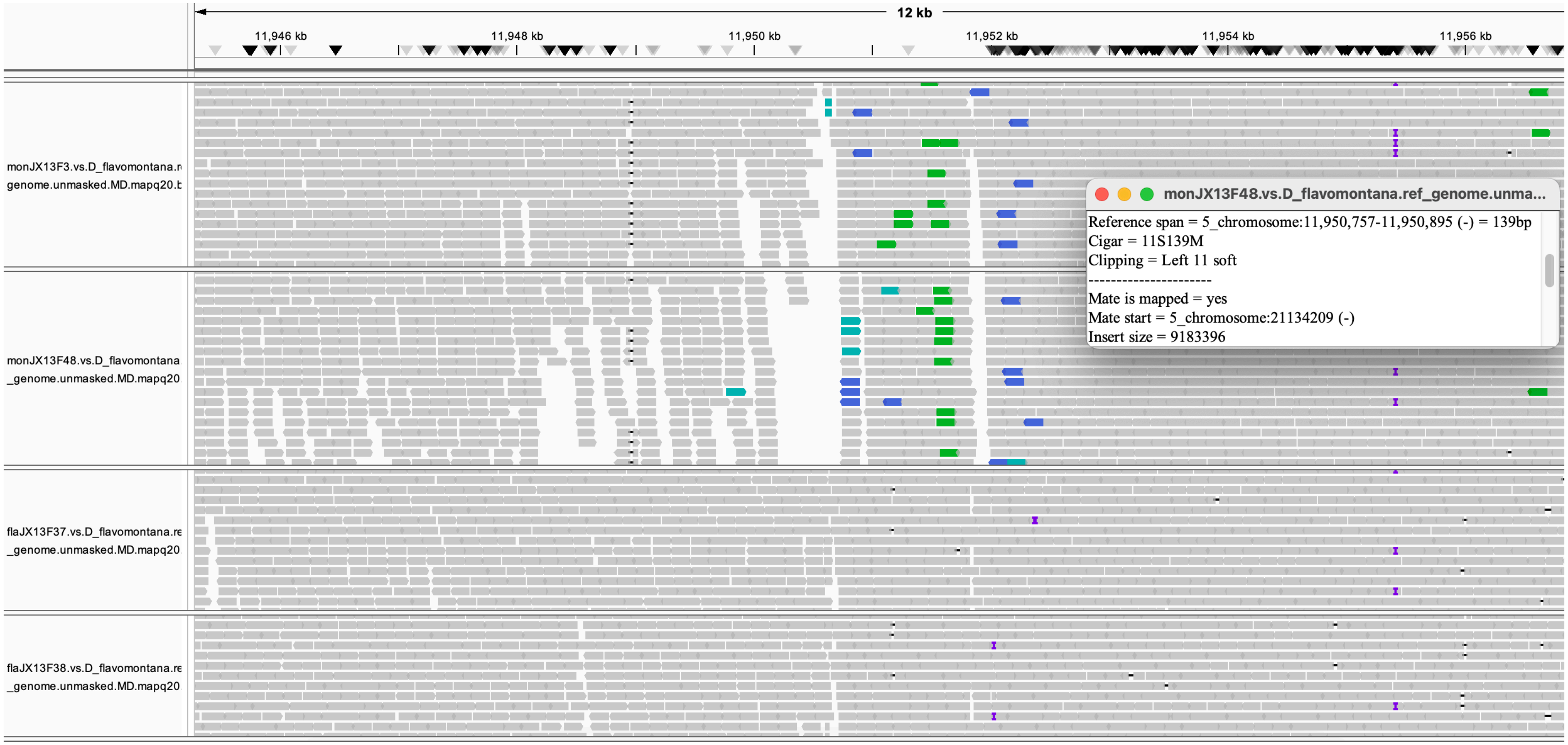
12 kb
11,946 kb
11,948 kb
11,950 kb
11,952 kb
11,954 kb
11,956 kb
monJX13F3.vs.D_flavomontana.r
genome.unmasked.MD.mapq20.b
monJX13F48.vs.D_flavomontana
_genome.unmasked.MD.mapq20
flaJX13F37.vs.D_flavomontana.re
_genome.unmasked.MD.mapq20
flaJX13F38.vs.D_flavomontana.re
_genome.unmasked.MD.mapq20
monJX13F48.vs.D_flavomontana.ref_genome.unma...
Reference span = 5_chromosome:11,950,757-11,950,895 (-) = 139bp
Cigar = 11S139M
Clipping = Left 11 soft
----------------------
Mate is mapped = yes
Mate start = 5_chromosome:21134209 (-)
Insert size = 9183396

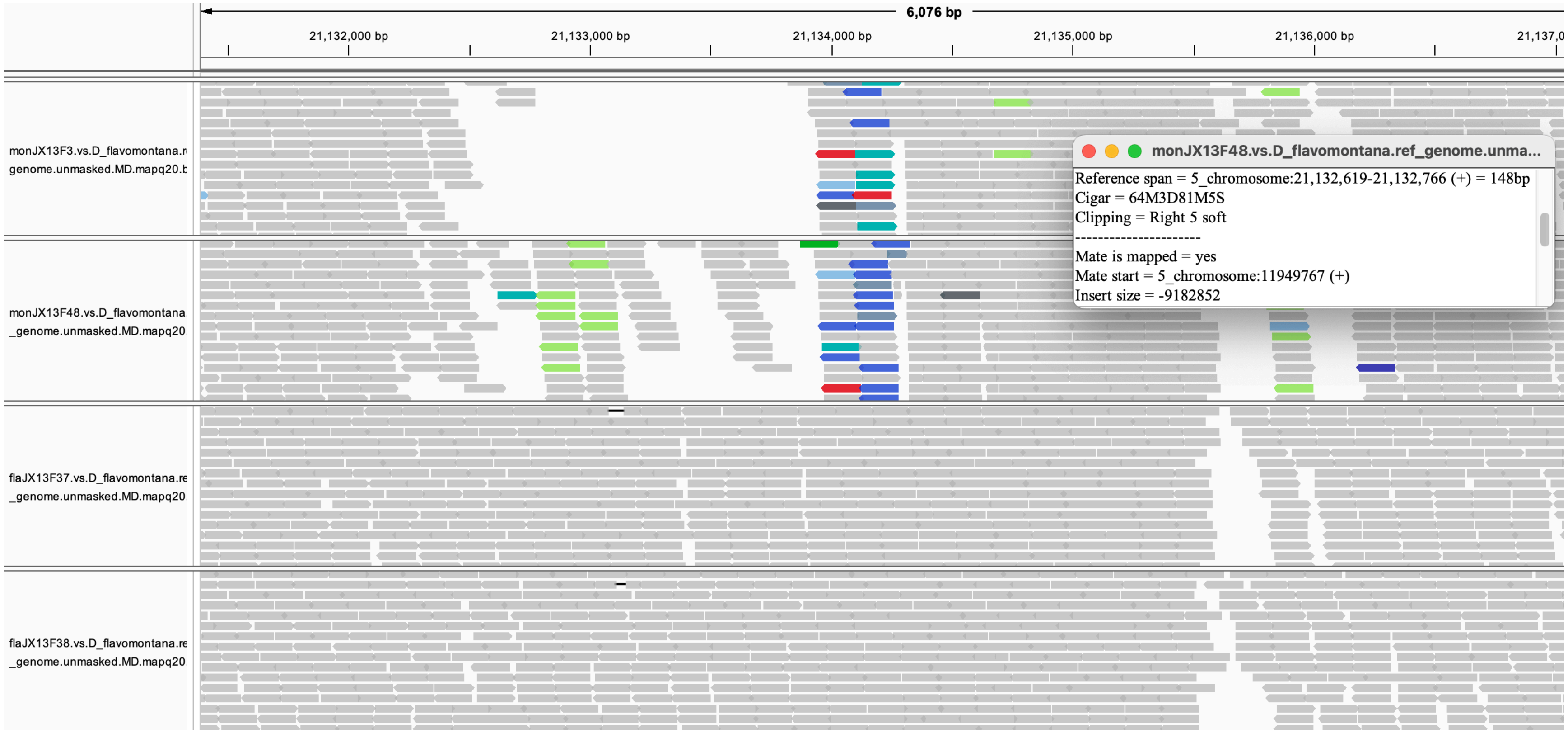
6,076 bp
21,132,000 bp
21,133,000 bp
21,134,000 bp
21,135,000 bp
21,136,000 bp
monJX13F3.vs.D_flavomontana.r
genome.unmasked.MD.mapq20.b
monJX13F48.vs.D_flavomontana
_genome.unmasked.MD.mapq20
flaJX13F37.vs.D_flavomontana.re
_genome.unmasked.MD.mapq20
flaJX13F38.vs.D_flavomontana.re
_genome.unmasked.MD.mapq20
monJX13F48.vs.D_flavomontana.ref_genome.unma...
Reference span = 5_chromosome:21,132,619-21,132,766 (+) = 148bp
Cigar = 64M3D81M5S
Clipping = Right 5 soft
Mate is mapped = yes
Mate start = 5_chromosome:11949767 (+)
Insert size = -9182852

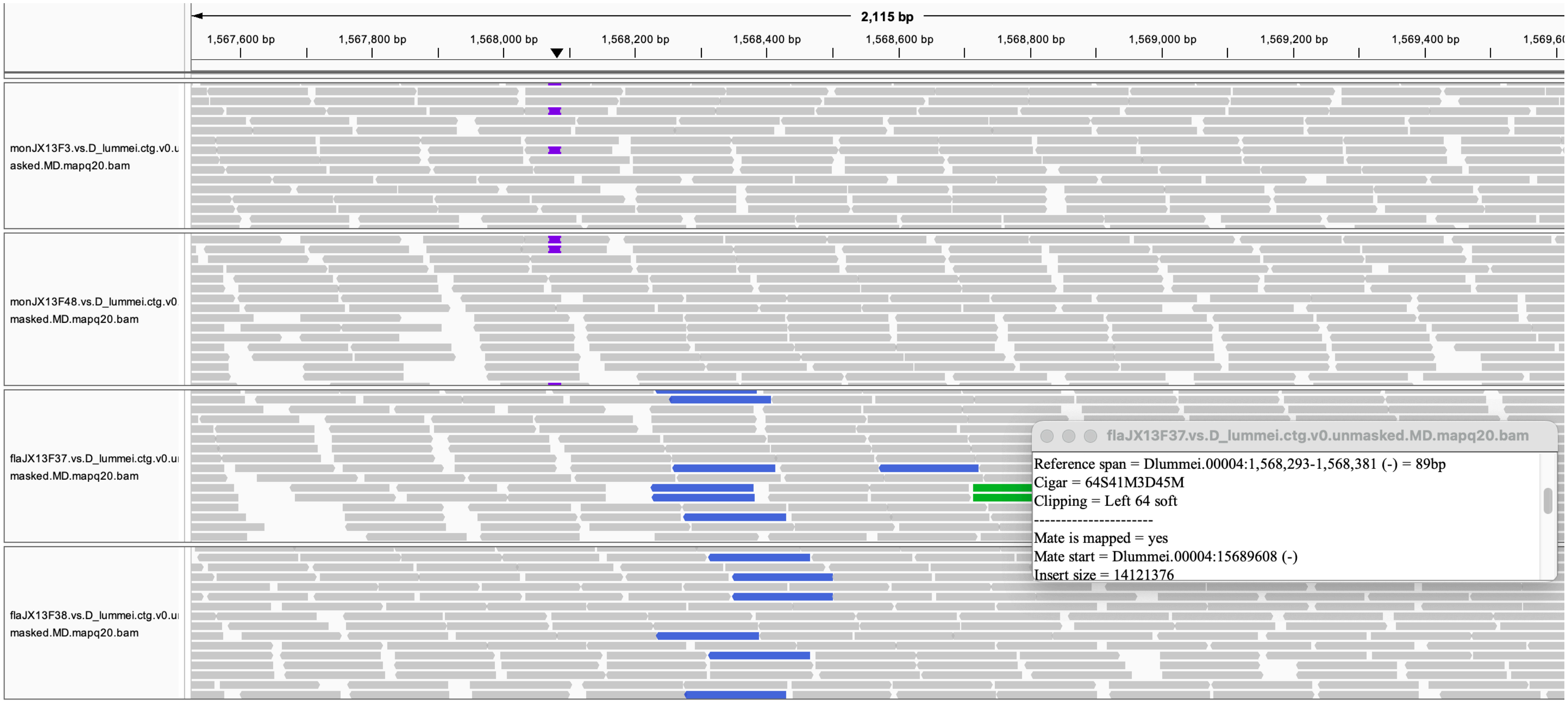
2,115 bp
1,567,600 bp
1,567,800 bp
1,568,000 bp
1,568,200 bp
1,568,400 bp
1,568,600 bp
1,568,800 bp
1,569,000 bp
1,569,200 bp
1,569,400 bp
monJX13F3.vs.D_lummei.ctg.v0.u asked.MD.mapq20.bam
monJX13F48.vs.D_lummei.ctg.v0 masked.MD.mapq20.bam
flaJX13F37.vs.D_lummei.ctg.v0.u masked.MD.mapq20.bam
flaJX13F38.vs.D_lummei.ctg.v0.u masked.MD.mapq20.bam
flaJX13F37.vs.D_lummei.ctg.v0.unmasked.MD.mapq20.bam
Reference span = Dlummei.00004:1,568,293-1,568,381 (-) = 89bp
Cigar = 64S41M3D45M
Clipping = Left 64 soft
----------------------
Mate is mapped = yes
Mate start = Dlummei.00004:15689608 (-)
Insert size = 14121376

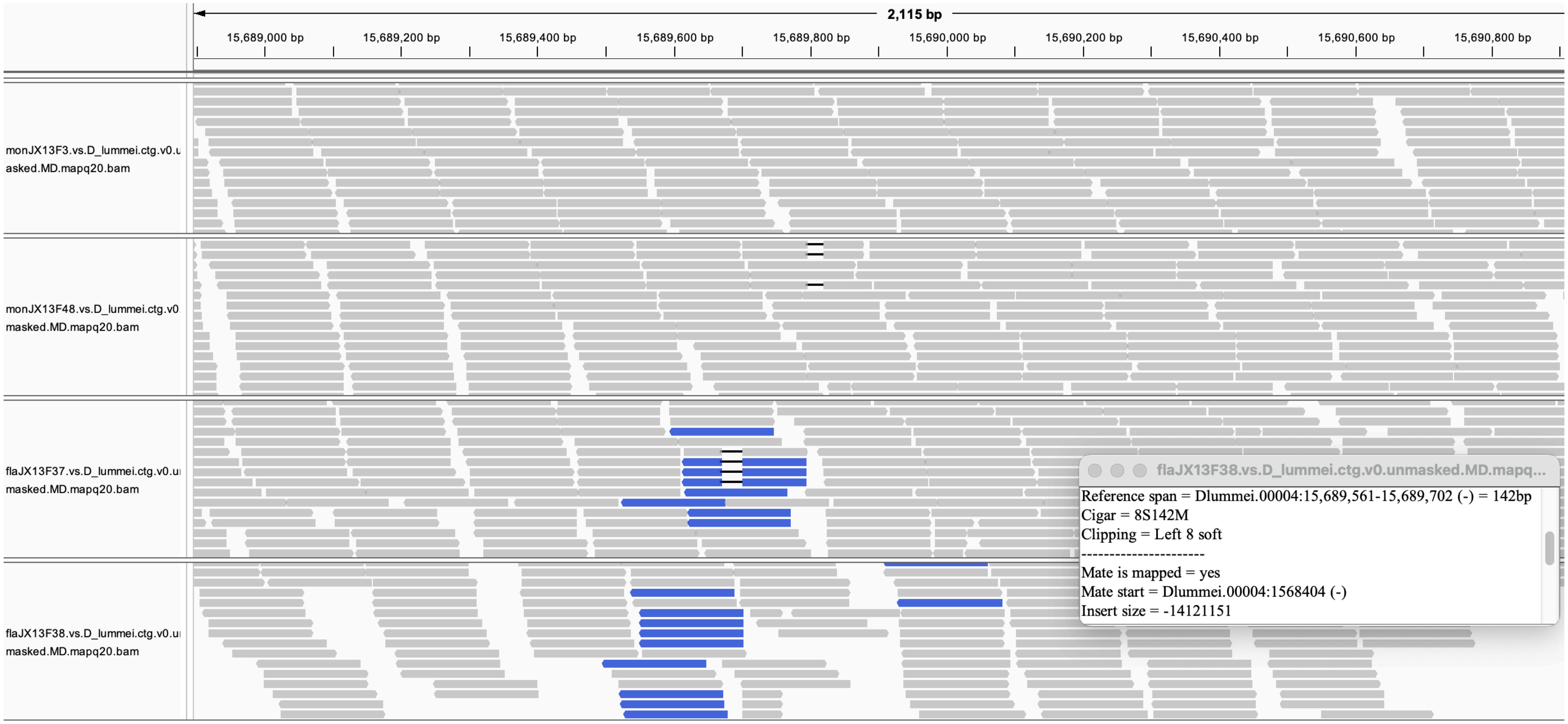

Figure S3. Inversion breakpoints of alternatively fixed inversions between *D. montana* and *D. flavomontana* pools. Illumina paired-end re-sequencing data of both species were mapped against *D. flavomontana* reference genome, and the data is illustrated with Integrative Genomics Viewer (IGV) [3]. Due to inversion differences between the species, *D. montana* reads are split, the insert size of its paired reads deviates from the expected (red reads), and paired reads are in reversed orientation (blue and turquoise reads). Note that not all X inversion breakpoints exhibit reversed orientation due to the complex overlap of these inversions (see Table S1). Since chromosome 4 of the *D. flavomontana* reference genome was scaffolded, the presence of the inversion on chromosome 4 was verified by mapping both species to the *Drosophila lummei* reference genome (breakpoints 1,568,294 and 15,689,984 in the *D. lummei* genome equal to 23,745,872 and 7,855,585 in the *D. flavomontana* genome, respectively; Table S1), as described in Poikela et al. [1]. The data underlying this figure can be found at doi.org/10.5281/zenodo.22277897.

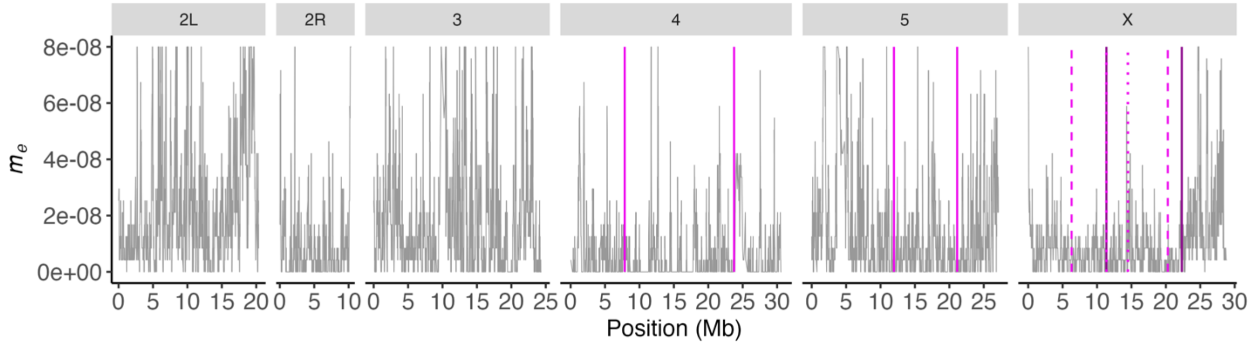


Figure S4. Variation in migration rate ($m_e$) across sliding windows between natural populations of *D. flavomontana* and *D. montana*. Vertical solid and dashed lines denote the breakpoints of the alternatively fixed chromosomal inversions between *D. flavomontana* and *D. montana*. The data underlying this figure can be found at doi.org/10.5281/zenodo.22277897.
